# Y-linked copy number polymorphism of target of rapamycin (TOR) is associated with sexual size dimorphism in seed beetles

**DOI:** 10.1101/2023.04.14.536871

**Authors:** Philipp Kaufmann, R. Axel W. Wiberg, Konstantinos Papachristos, Douglas G. Scofield, Christian Tellgren-Roth, Elina Immonen

**Affiliations:** Department of Ecology and Genetics (Evolutionary Biology program), Uppsala University, Norbyvägen 18D, 75234 Uppsala, Sweden; Department of Cell and Molecular Biology, Molecular Evolution, Uppsala University, Sweden; Uppsala Multidisciplinary Center for Advanced Computational Science, Uppsala University, Uppsala, Sweden; National Genomics Infrastructure, Uppsala Genome Center, SciLifeLab, BioMedical Centre, Uppsala University, 751 08, Uppsala Sweden; Ecology Division, Department of Zoology, Stockholm University, Svante Arrhenius väg 18B, 106 91, Stockholm, Sweden

**Keywords:** Y chromosome, Y polymorphism, sex chromosome, sexual dimorphism, Callosobruchus maculatus, sexual conflict, Target of rapamycin, TOR

## Abstract

The Y chromosome is theorized to facilitate evolution of sexual dimorphism by accumulating sexually antagonistic loci, but empirical support is scarce. Due to the lack of recombination Y chromosomes are prone to degenerative processes, which poses a constraint on their adaptive potential. Yet, in the seed beetle *Callosobruchus maculatus* segregating Y linked variation affects male body size and thereby sexual size dimorphism (SSD). Here we assemble *C. maculatus* sex chromosome sequences and identify molecular differences associated with Y-linked SSD variation. The assembled Y chromosome is largely euchromatic and contains over 400 genes, many of which are ampliconic with a mixed autosomal and X chromosome ancestry. Functional annotation suggests that the Y chromosome plays important roles in males beyond primary reproductive functions. Crucially, we find that, besides an autosomal copy of the gene *target of rapamycin* (*TOR*), males carry an additional *TOR* copy on the Y chromosome. TOR is a conserved regulator of growth across taxa, and our results suggest that a Y-linked *TOR* provides a male specific opportunity to alter body size. A comparison of Y haplotypes associated with male size difference uncovers a copy number variation for *TOR*, where the haplotype associated with decreased male size, and thereby increased sexual dimorphism, has two additional *TOR* copies. This suggests that sexual conflict over growth has been mitigated by autosome to Y translocation of *TOR* followed by gene duplications. Our results reveal that despite of suppressed recombination, the Y chromosome can harbour adaptive potential as a male-limited supergene.

## Introduction

Driven by differences in reproductive strategies^1, 2^ males and females commonly experience sexually antagonistic selection on traits present in both sexes ^e.g.3, 4^. When the sexes share genetic variation in homologous traits^5^, an allele beneficial to females may be deleterious for males and *vice versa*. This genetic dependency can hinder sex-specific adaptations, causing intralocus sexual conflict^6^. The Y chromosome is limited to males, and linkage of sexually antagonistic loci on the Y would represent a straightforward solution to this genetic conflict^7–10^. A male limited pathway to alter the expression of a sexually antagonistic trait disconnects the genetic basis between the sexes and can reduce gender load by enabling males but also females to reach their fitness optima.

Yet, the role of Y chromosomes in the evolution of sexual dimorphism has historically been neglected^7^. This is because the Y typically degenerates quickly after the loss of recombination with the X, which is expected to constrain its adaptive potential^11^. The Y chromosome degenerates due to reduced efficacy of selection and mutation accumulation (reviewed in^12^), selective haploidisation driven by regulatory evolution^13^ or a combination thereof. As a consequence of degeneration, the Y chromosomes are often heteromorphic from the X, void of genes and rich in repetitive elements. Y chromosome sequences are therefore difficult to assemble, and has been done only in a handful of mammalian species such as human^14^, other primates^15, 16^, mouse^17^ and bull^18^. There are also a few examples of near complete assemblies such as for malaria mosquitos^19^, some fish^20, 21^ and neo-Y in *Drosophila miranda*^22^. Genes remaining on degenerate Y chromosomes are frequently translocated from the autosomes, allowing their male specialisation. Y linked genes show typically testis specific expression ^e.g.14, 16, 17, 20^, which suggests they are associated with male primary reproductive functions. But recently, heteromorphic Y chromosomes have gotten more attention in regards to sexually dimorphic, non-reproductive traits in humans^23^, drosophila^24^, colour and behavioural variation in fish^21^, and body size in seed beetles^25^.

The order of Coleoptera (beetles) is the most species rich group of animals on the earth. Yet only a handful of beetle species have been sequenced thus far^26^, and their Y chromosomes remain largely uncharted. Coleoptera has XY sex determination but the exact mechanism is unknown. A study of karyotypes of over 4000 beetle species shows occasional loss of Y chromosomes^27^, suggesting that Y is not essential for sex determination^28^. Interestingly, most studied species in the largest Coleopteran suborder Polyphaga do not have an obligate XY chiasmata formation, and therefore lack a pseudo autosomal region (PAR) and XY recombination altogether^27^. One such species is *Callosobruchus maculatus* seed beetles that harbours a heteromorphic Y chromosome estimated to represent less than 2% of the genome^29^. Based on the karyotype, the *C. maculatus* Y is euchromatic despite its small size^30^.

We recently discovered segregating Y-linked genetic variance that alters male body size^25^. Female and male *C. maculatus* have different fitness optima for body size^31–33^, which is closely connected to many other life history traits^34^. A combination of quantitative genetic analysis, artificial selection and isolating the effect of Y linked genetic variance by introgressing the putative Y haplotypes onto a common genetic background, revealed two phenotypically distinct male morphs associated with different Y haplotypes (Y_L_ and Y_S_ for large and small size, respectively). We could demonstrate that directional selection on males can deplete Y haplotype variation quickly^35^, and that carrying either one of the Y haplotype alters the male size (weight), and consequently the level sexual size dimorphism, by 30%^25^. Body size is a classic quantitative trait, and while quantitative genetic evidence suggests that even the autosomal genetic architecture of body size in *C. maculatus* consist of a combination of few major effect loci and many small effect loci^35^, finding that a substantial part of its architecture in males is controlled by a non-recombining ‘supergene’ is surprising. But a Y-linked element has also been implicated in male body size variation in humans^36^ and fish^21, 37^ suggesting that Y-linkage may be a common way to mitigate sexual conflict over growth across diverse taxa.

In this study we assembled previously un-characterised X and Y chromosome sequences of *C. maculatus*, and identified molecular differences between the two Y haplotypes (Y_S_ & Y_L)_ associated with small or large male body size and with major effects on sexual size dimorphism, shedding light on the underlying molecular mechanisms. To do this we took advantage of comparing genomes of the Y_S_ and Y_L_ introgression lines^25^ (hereafter referred to as S and L, respectively) that share inbred autosomal and X chromosomes and only differ in their respective Y haplotype. Here, we first identified non-recombining X and Y contigs in both S and L genomes, by comparing sequence coverage difference between males and females, and verified male specificity of the longest Y contig (8.4Mb) with PCR. To further confirm the identity of the Y contigs and characterise Y variation, we compared the genomes of S and L lines that share variants in all other chromosomes except on the Y. We identified and functionally characterised protein coding genes and repeat structure of the Y contigs, analysed the origins of the Y genes as X gametologs or autosomal paralogs, studied gene duplications within the Y to identify putative ampliconic genes, and examined expression of Y-linked genes.

## Results

### Genome assembly

First, we separately assembled the genomes of the S & L lines, and annotated the S genome (carrying the ancestrally more frequent Y_S_ haplotype), which was subsequently used as the reference genome in this study. The assembled S genome has 938 contigs that yield a total genome length of 1.246 Gb. The N50 value of the assembly is 9.45 Mb with the longest contig being 37.4 Mb in length, the L50 is 38 and the BUSCO completeness scores over 98% (insecta_odb10: complete: 98.1% [single copy: 87.6%, double copy: 10.5%], fragmented: 0.2%, missing: 1.7%, n=1367), demonstrating that the assembly is of high quality and further improves the previously published genome for this species (*C. maculatus* reference genome; N50 of 0.15 Mb and total genome size of 1.01 Gb^29^). 597 contigs were shorter than 100 kb (consisting of 24.9 Mb length, 2.00% of the total S assembly length) and were not considered in the downstream sex chromosome identification analysis because their chromosome type (autosome, X or Y) could not be determined with confidence due to their short length. The L line assembly is also of high quality with 1323 contigs and a total length of 1.224 Gb, the longest contig being 30.2 Mb in length, with an N50 of 9.86 Mb, L50 of 38 and a BUSCO completeness score of over 97% (insecta_odb10: complete: 97.8% [single copy: 87.5%, double copy: 10.3%], fragmented: 0.3%, missing: 1.9%, n=1367). 945 contigs were shorter than 100 kb (consisting of 41.0 Mb, 3.35% of the total L assembly length).

After repeat-masking, 72.1% of the reference genome was soft-masked, of which 21.1% was identified as retroelements (primarily LINEs: 17.8%) and 21.0% as DNA transposons. A total of 24.8% interspersed repeats remained unclassified. Various low-complexity repeats formed the remainder of the soft-masked content. The final set of annotated gene models include 35,865 genes (68% increase compared to the original assembly^29^), 39,983 transcripts (3451 two-transcript gene models and 297 with more than two transcripts; 14% increase in the total number of transcripts^29^). A total of 25,651 transcripts received functional annotation.

### Identification of Y and X contigs

We performed a coverage comparison analysis (with SATC^38^) using Illumina short-read sequencing data from samples of both sexes^29^ to identify novel sex chromosome sequences. SATC correctly identified the sexes of the samples. In the S genome assembly four Y (total of 10.1 Mb) and eight X contigs (total of 58.6 Mb) were detected based on significant coverage differences, while in the slightly more fragmented L genome we identified five Y contigs (total of 4.89 Mb) and ten X contigs (total of 64.2 Mb) (Table S1-S2). Importantly, the identified Y contigs from both assemblies map to each other, demonstrating that SATC identified homologous sequences in both assemblies (Fig. S1-S2). The same contigs were identified whether using unfiltered data or when using repeat masked contigs, with minor exceptions (see Table S1-S2 and Fig. S1-S2). Note that the SATC analysis identified several additional contigs to show significant coverage difference between the sexes (see online Table O1), however, the relative coverage difference in these cases was below 10% and we therefore took a more conservative approach and only considered sex chromosome contigs above this threshold in our downstream analysis. It is possible however that these (or other unidentified) contigs are still sex linked but less diverged between X and Y.

Gene ontology (GO) enrichment of Y linked transcripts, as compared to all identified sex-linked transcripts, shows that Y is functionally different from the X (Fig. 1). The significantly enriched processes on the Y include cell proliferation, regulation of development, cell death and apoptosis, response to stress/external stimulus such as response to starvation, immune response, RNA processing and regulation of posttranscriptional gene expression as well as protein modification (ubiquitination), and various metabolic processes (Fig. 1 & online Table O2). GO enrichment of Y transcripts for molecular function are presented in Fig. S7.

**Fig. 1:**
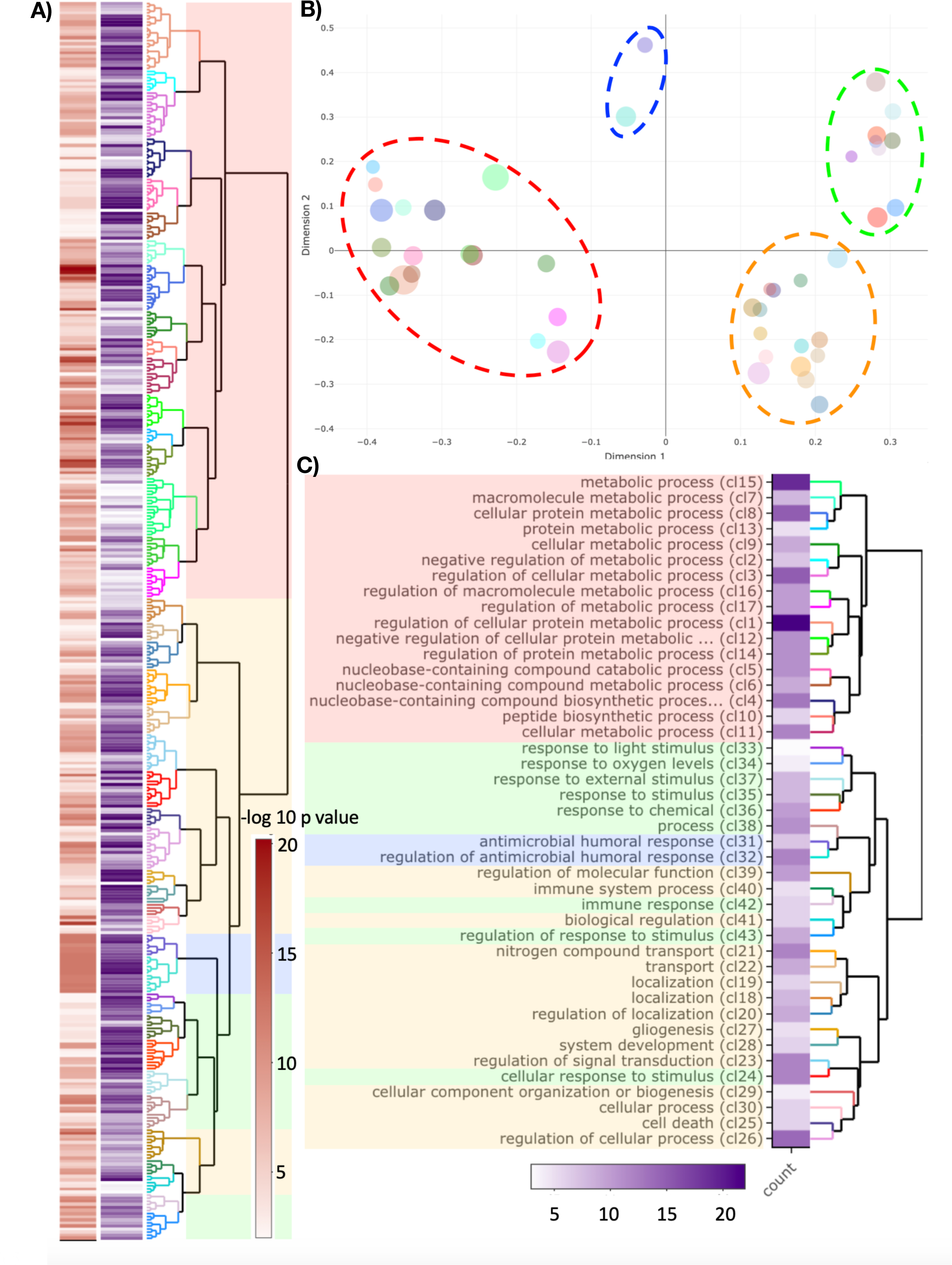
Gene ontology enrichment for Y-linked genes with ‘biological process’ annotations. **A)** Dendrogram of GO terms based on Wang’s semantic similarity distance, heatmap (red) indicates statistical significance as −log_10_ p-values (i.e. higher −log_10_ p-values have higher statistical support) and information content (purple). **B)** Multi-Dimensional Scaling (MDS) plot based on Best-Match Average (BMA) distance, representing the proximities of dendrogram clusters in (**A**). Dot size indicates the number of GO terms within each cluster. We highlight the four major functional groups with dashed ellipsoids (red ≅ metabolic processes; blue ≅ antimicrobial response; green ≅ response to stimulus; yellow ≅ development, cell organisation, growth & cell apoptosis) **C)** Dendrogram representation of clusters from (**B**) with GO term description of the first common GO ancestor and heatmap for the number of GO terms within each cluster.

The identified Y contigs, in either the Y_S_ or the Y_L_ haplotype, do not seem to show a higher number of repeats per length, nor a higher percentage of repeat content, compared to the X or autosomal contigs (Fig. S4-S5). However, the composition of repeat content on the Y is somewhat unique, where repeats identified as DNA/Maverick and LINE/Penelope are overrepresented, as a proportion of all repeat elements, on the Y compared to the X or autosomal contigs (Fig. S6).

### Gametologs, paralogs and ampliconic genes

We detected 437 transcripts on the Y_S,_ of which 202 transcripts are ampliconic, and have between 1 and 13 additional nearly identical copies on the Y (>99.9 % nucleotide similarity), forming 67 ampliconic groups. Hence, we identified 302 unique Y-linked transcripts, of which 235 are non-ampliconic transcripts and 67 form ampliconic groups. 424 transcripts have at least one gametolog or paralog in the genome (when searching for Y protein sequences against all *C. maculatus* proteins, using blastp^39^ (>50% query coverage) and filtering for >80% sequence identity and E-value threshold = 1e-20). With these criteria we detected in total 281 unique autosomal paralogs, 214 unique X-linked gametologs and 359 unique Y-linked homologous transcripts (Fig. S9). When lowering the sequence similarity threshold, we find homologs even for the remaining 12 Y-linked transcripts, although the best hit protein sequence similarity drops to below 40% for some of them. To identify autosomal or X ancestry for each Y transcript, we categorized them as exclusively autosomal paralogs (n=157), exclusively gametologs on the X (n=99) or homologs on both (n=73) (Fig. 2), while excluding genes that have homologs on uncategorized contigs (<100 kb, n=51 genes). Y shared significantly more exclusive X gametologs than exclusive autosomal paralogs, when accounting for size difference (two-tailed Fisher’s exact test: 95%CI (9.56, 16.07), p < 0.0001) or difference in the overall number of transcripts (two-tailed Fisher’s exact test: 95%CI (8.32, 14.13), p < 0.0001). Gametologs that have been maintained on the Y are functionally enriched for RNA processing, particularly genes involved in RNA splicing, development as well as metabolic processes. Y transcripts with paralogs on the autosomes are functionally enriched for response to stimulus, developmental processes, protein translation and post-translation modification (Fig. S8). Interestingly, this elevated sequence similarity between the X and the Y is not reflected in pronounced sequence synteny blocks between X and Y contigs, neither in nucleotide sequence nor gametolog synteny (Fig. S10).

**Fig. 2:**
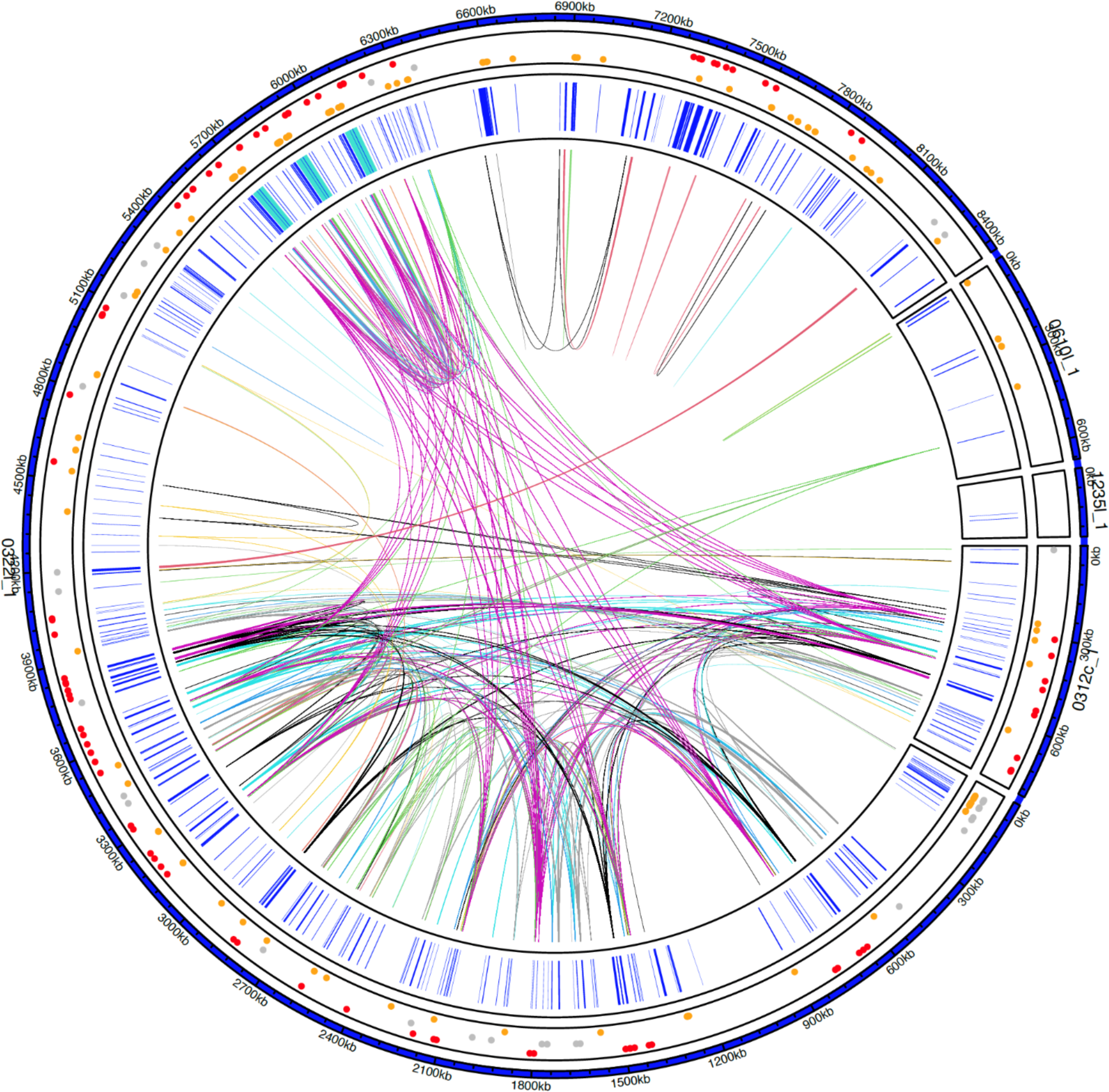
Overview of Y genes. Each amplicon group (i.e. genes with >99.9% nucleotide sequence identity and >95% query coverage) is highlighted with a genomic link. 219 out of a total of 437 Y genes have at least one additional copy on Y. Gene positions of all Y-linked genes are shown in inner track in blue, regions containing *TOR* are highlighted in turquoise (see Fig. 4A for more details). The outer track indicates whether a gene has exclusively gametologs on the X (red, n=99), paralogs on the autosomes (yellow, n=157) or both (grey, n=73), dots are scattered by homolog type.

### Characterisation of the Y variation associated with the body size difference

#### Variant calling

The patterns of shared (hereafter SNP) and fixed (hereafter fixed SNV) single nucleotide variants in the S & L genomes are well aligned with the expectations considering how the lines were created, and further confirm the identity of the detected X and Y sequences. The majority of single nucleotide variants (2,823,154) are shared polymorphisms in both genomes (2,812,361 SNPs, 99.6%) and there are only few fixed SNV differences between the two introgression lines (10,793, 0.38%) (Table 1). Autosomal contigs have significantly more shared SNP/bp than contigs identified as the sex chromosomes (two-tailed Fisher’s exact test: 95%CI (28.8, 30.5), p < 0.0001 and 95%CI (83.2, 109.6), p < 0.0001, for the X and Y respectively) (Fig. S11), and the X contigs have significantly more shared SNP/bp than the Y contigs (two-tailed Fisher’s exact test: 95%CI (2.80, 3.71), p < 0.0001). In contrast, the Y contigs have significantly more fixed SNV differences/bp than autosomal contigs (two-tailed Fisher’s exact test: 95%CI (43.4, 47.3), p < 0.0001), and there are no fixed SNV differences on the X contigs (Fig. S12).

**Table 1:** The number of SNP and fixed differences between the two Y introgression lines split by chromosome type. Note that contigs shorter than 100 kb are not categorized as either Y, X or A.

We identified a total of 137 genes with fixed SNV differences within the gene region or in close proximity to genes (i.e. ±2 kb as an approximation of cis-regulatory up and downstream area) between the two Y-haplotypes (Fig. 3). For 12 out of these 137 genes, we could identify unique *D. melanogaster* orthologs and annotate their function via FlyBase^40^ including DNA/RNA binding, mRNA splicing, regulation of cell proliferation and protein ubiquitination (Table S3, further details in online Table O3).

**Fig. 3:**
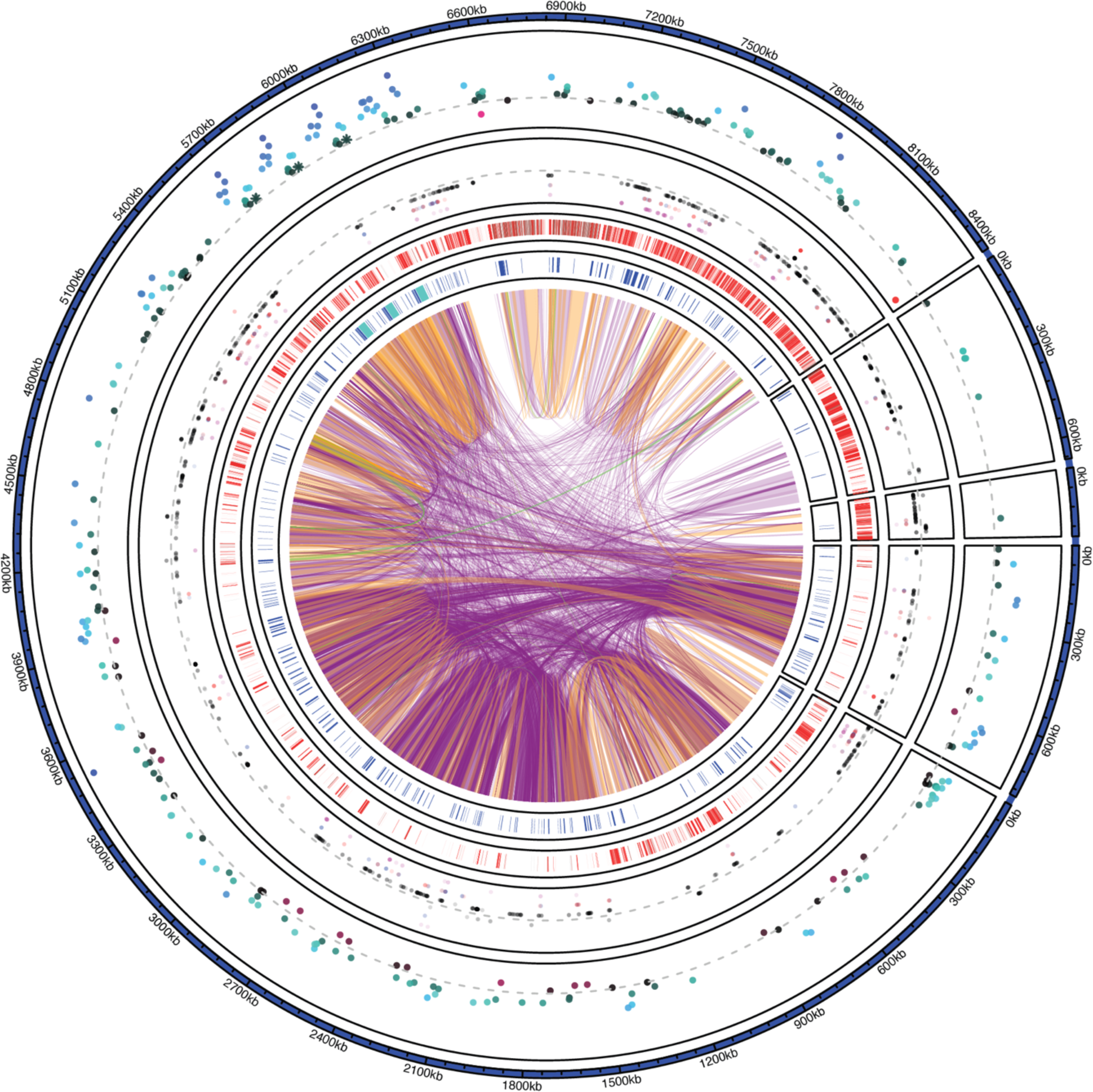
Overview of Y contigs in the Y_S_ genome. Color coded genomic links show high level of nucleotide sequence similarity among and within Y contigs (purple > 99%, orange > 99.9%, green >99.99% nucleotide similarity matches > 1 kb in size). The inner track shows the position of annotated genes in blue, with the *TOR* regions highlighted in turquoise (Fig. 4). The second inner track highlights areas of low mapping coverage between the two Y haplotypes: areas shown in red indicate coverage lower than 17x (i.e. regions that may lack coverage for reliable variant calling via DeepVariant). The third track shows the position of single nucleotide variants (SNV), single nucleotide polymorphisms (SNPs) (outside of the dashed grey line) and fixed SNV (inside the dashed grey line). SNV in black (closest to the dashed grey line) are outside of gene regions, SNV in magenta (farthest from the dashed grey line) are within gene regions. SNV in blue and red are in 2 kb upstream or 2 kb downstream (proxy for cis-regulatory region of a gene) of a gene. The outer track shows differential gene expression between males and females. Genes in blue, towards the outside are male biased, genes in red, towards the center, are female biased (grey dashed line is shown as a reference to indicate no difference in gene expression). *TOR* gene expressions are highlighted as asterisks and are significantly male biased.

#### Y-linked TOR amplicon

One of the annotated Y-linked genes indicated strong homology to the gene *target of rapamycin (TOR),* a highly conserved member of the IIS/TOR pathway^41^. Mapping a consensus *TOR* protein to our assembly via exonerate^42^ identified one autosomal *TOR* gene, detected in both Y_S_ and Y_L_ genomes (see supplementary methods ‘genome annotation’ for full details). In addition, there is one gene on the opposite strand that matches to adenosine deaminase 2 in several taxa (also involved in cell proliferation).

We further discovered three consecutive copies of the Y *TOR* on the Y_S_ haplotype that causes the small body size morph in males (Fig. 4A), but there is only a single Y-linked *TOR* in the Y_L_ haplotype associated with the large body size morph in males (Fig. 4B), revealing Y-linked copy number variation (CNV) of the *TOR* region. Aligning Y_S_ and Y_L_ contigs (Fig. S13) shows that the *TOR* region CNV is located in the middle of an otherwise continuous alignment between two haplotypes, showing that CNV is not an artefact caused by a broken Y_L_ haplotype contig.

**Fig. 4:**
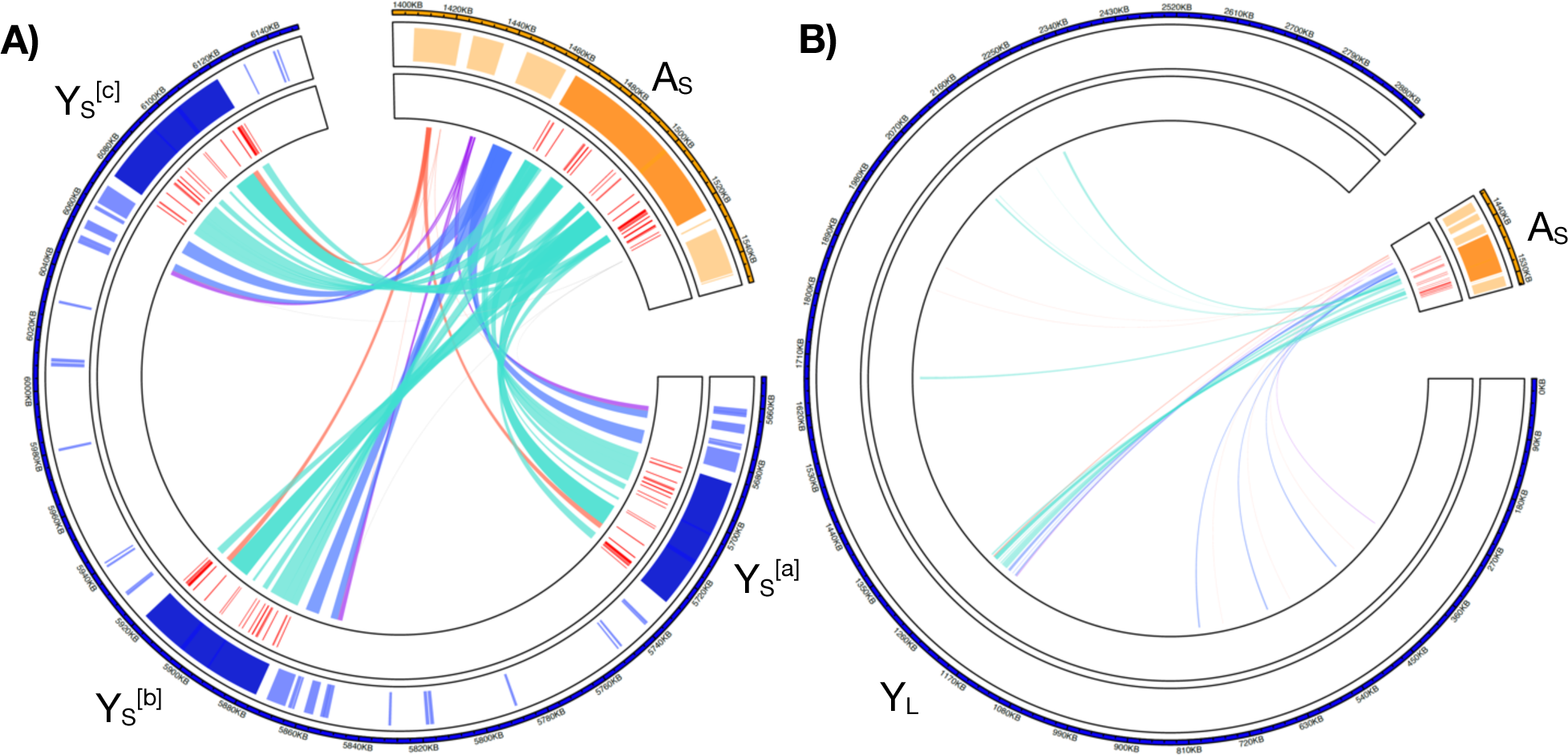
Zoom into the *TOR* region on the Y (blue) and on the autosome (yellow). The inner track shows the position of identified exons in red. Note that all Y linked *TOR* copies lack the exons 1-5 (from the 5’ end), have one partial exon 6, all the other *TOR* exons (7-25) are present and complete. **A)** Y_S_ haplotype, showing the three *TOR* regions (denoted with ^[a], [b]^ & ^[c]^). Note that we here only present the relevant region of the whole utg0003221_l Y_S_ contig (blue, see highlighted region in Fig. 2&3). **B)** Y_L_ contig containing the *TOR* region in the Y_L_ haplotype (blue, showing the whole contig) and the autosomal *TOR* region (yellow, from the annotated Y_S_ assembly). In the Y_L_ haplotype we identify only one Y-linked copy.

A closer comparison of the autosomal and the Y linked *TOR* region reveals that all Y-linked *TOR* copies (in both Y_S_ and Y_L_ haplotypes) lack the initial five 5’ CDS compared to the autosomal *TOR*. A maximum likelihood tree (Fig 5A) of concatenated CDS comparing all *TOR* regions (i.e. the autosomal *TOR* of each assembly (A_S_ & A_L)_, three Y-linked Y_S_ *TOR* copies and one Y-linked Y_L_ *TOR*) shows the following; First, the autosomal A_S_ & A_L_ *TOR* regions are isogenic, as expected. Second, all Y-linked *TOR* sequences cluster together with high bootstrap confidence, indicating that the *TOR* transposition from the autosome to the Y predates the two Y haplotypes. Also, DeepVariant SNP calling did not detect any SNV differences within the *TOR* region between the Y_S_ and Y_L_ haplotypes, which might be due to lower coverage when mapping the two Y haplotypes against each other (Fig. 3). We find high confidence that Y s^[a]^ & Y s^[b]^ *TOR* copies are most similar to each other but the remaining clustering of Y-linked *TOR* sequences has low support. Importantly, while the exons align with high similarity across all *TOR* regions, non-coding sequences have diverged more and show lower sequence similarities (Fig. 5B). This is particularly apparent when comparing autosomal and Y-linked *TOR* regions, where non-coding sequences frequently do not align, indicating structural differences between them (Fig. 5B, Fig. S13).

**Fig. 5:**
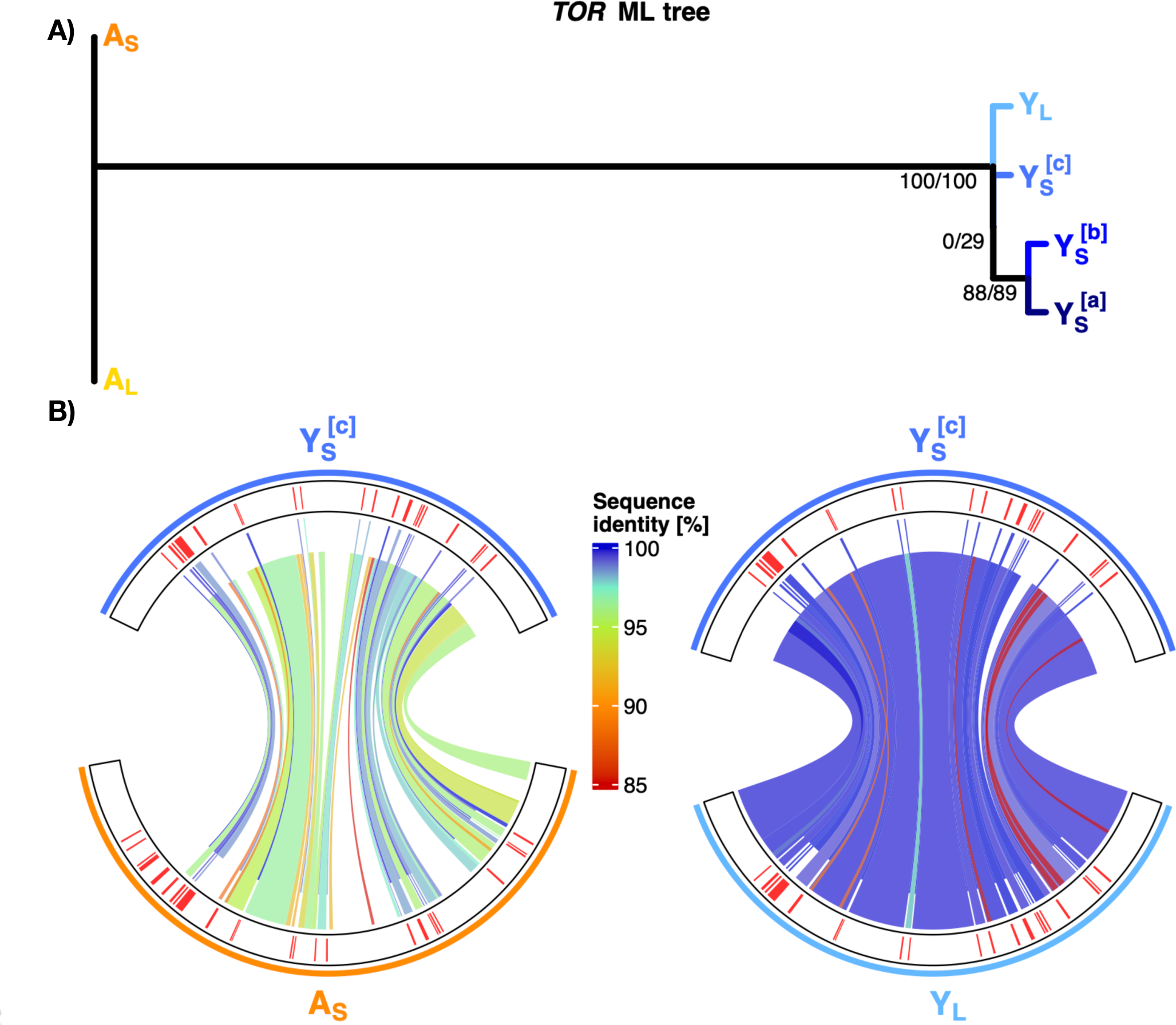
*TOR* region genealogy and comparison. **A)** Maximum likelihood gene tree of exonic *TOR* sequences. Note that the A_S_ and A_L_ (highlighted in orange shades) have identical exonic sequences. High bootstrap values (100/100) indicate that Y-linked *TOR* (highlighted in blue shades) are more similar to each other than to the autosomal *TOR* and that Y s^[a]^ & Y s^[b]^ are sister sequences to each other (88/89). However, the clustering of the remaining Y-linked *TOR* sequences has low support (0/29). **B)** Nucleotide alignment with Mummer of exonic and non-coding regions separately. The outer track shows the *TOR* exon positions in red. Genomic links show nucleotide similarity, exonic links are shifted upwards, while non-coding alignments are shifted down to guide easier distinction between the two regions. Fully factorial pairwise comparison of all *TOR* regions is presented in Fig. S13. Left side: Comparison of the autosomal and the Y_S[c]_ *TOR* region. While there are structural differences (gaps) and low sequence identity matches in the non-coding regions, the exonic regions seem conserved between the autosomal and the Y_S_ *TOR* copy. Note that all Y *TOR* regions lack the exon 1-5 (from the 5’ end) and exon 6 is only partially present. Exon 7-25 are present in all Y *TOR* regions. Right side: Comparison of Y s^[c]^ *TOR* and the *TOR* region on the other Y haplotype (Y). We find high nucleotide similarity in both non-coding and exonic regions alike, with one structural difference in a non-coding region.

In addition to the copy number variation of the *TOR* locus, aligning the Y_S_ and Y_L_ haplotypes to each other suggests that there are further structural differences between the haplotypes (Fig. S14). Moreover, there are similarities between the classes and distribution of repetitive elements between the autosomal and the different copies of the Y-linked *TOR* region, indicating homology also at the level of *TOR*-associated repetitive elements (Fig. S15).

## Discussion

Here we assembled two new *C. maculatus* genomes. We successfully identified 10 Mb of the Y chromosome, using a set of complementary methods, and analysed the gene content of the Y sequences. We discovered that despite having lost most of its sequence since divergence from the X, the *C. maculatus* Y is rich in genes, many of which are expressed, with diverse functions from metabolism, development and stimulus response to regulation of gene expression and translation. This is in line with karyotype information suggesting that the Y is mostly euchromatic^30^, in spite of not recombining. We characterized molecular differences between the two Y haplotypes that were previously inferred from patterns of male limited inheritance of body size^25^. We identified a Y-linked copy of a functionally well conserved autosomal growth factor gene *target of rapamycin* (*TOR*), highlighting the potential for male specific growth regulation via the Y chromosome. Furthermore, we detected copy number variation of *TOR* between the two Y haplotypes underlying body size variation. Together our results indicate a central role of the Y chromosome in the evolution of sexual dimorphism in *C. maculatus* via male-specific evolution of *TOR*.

### XY identification

Degenerate sex chromosomes are notoriously difficult to assemble^43^ and in *C. maculatus* the entire genome has a high repeat content^29^. Despite these challenges, long read sequencing yielded high quality and contiguous genome assemblies for both Y introgression lines that allowed us to identify large parts of both the X and the Y chromosomes, greatly extending and curating previously identified sex chromosome-linked portions of the genome. Flow cytometry experiments in *C. maculatus* have previously given estimates of ∼18 and 93Mb for the size of Y and X chromosomes, respectively, although there is great uncertainty in the size estimate for the Y^44^. We could assemble four Y contigs yielding 10.1 Mb, ∼56% of the current length estimate. By comparison, in *D. melanogaster* ∼10% of the Y has been assembled thus far^45^, and to our knowledge no Y assemblies exist yet for other beetle species. Additionally, we assembled a total of 58.6 Mb of the X chromosome, corresponding to approximately 63% of the estimated size. Sex chromosome contig identification depends on the quality and length of contigs, accuracy and completeness of the genome, and, importantly, on the state of differentiation between the X and Y chromosomes. Here we rely on a suite of complementary methods including coverage difference between male and female reads, as well as genomic patterns of single nucleotide variants (SNVs) between our Y introgression lines that share the genome apart from the non-recombining parts of the Y chromosome. Mapping of previously described putative X and Y contigs^29^ also co-localize to our identified X and Y contigs in our assemblies, as expected. Furthermore, transcript expression patterns across the identified contig groups are in line with the unequal distribution of sex chromosomes between the sexes and show exclusive expression, or significantly higher male bias, of Y linked transcripts, and significantly less male bias for X-linked transcripts, as compared to the autosomal background. A large proportion of Y-linked transcripts show high sequence similarly with X-linked transcripts, in line with their common ancestry. Crucially, we confirmed the largest identified Y contig (8.4Mb) – carrying the identified Y linked *TOR* region – via male limited PCR amplification. The designed Y-specific primers work for both identified Y haplotypes and enables molecular sexing, a method that has thus far been lacking for *C. maculatus*.

### Gene content on the Y

We find 437 Y-linked transcripts (417 genes) and the identified Y contigs show high gene density, which is similar to the gene rich Y chromosomes characterized in mouse^17^ and bull^18^ but in contrast with the general expectation that Y-chromosomes should be low in gene content^7^, and other characterized insect Y chromosomes (*Drosophila*^46^ and mosquito^19^) that harbour only few genes. Genes retained on the Y, despite its degeneration, can broadly be categorized into two classes, genes that are dosage sensitive X gametologs^47, 48^ and genes that are male beneficial, which may have been recruited to the Y via autosomal translocations. Many (often ampliconic) Y genes show testis limited expression and are likely central to the male reproductive function^14, 15, 17, 20, 49, 50^. However, testis specific expression of highly amplified genes may also be indicative of meiotic drive^17, 18^, in which case the involved genes need not be male beneficial but Y beneficial instead. Y is predicted to accumulate sexually antagonistic genes only beneficial to males^8^, but there is still little direct evidence to support Y-linkage of traits that are also present in females. Metabolic rate, body size, longevity^31, 33, 51^, immunity and gene expression^29, 52, 53^ have all been implicated to be under sexually antagonistic selection in *C. maculatus.* We find functional enrichment of Y-linked genes that reflects these phenotypes remarkably well, including metabolic processes, immune response, response to stimulus and development, cell organisation, growth & cell apoptosis (Fig. 1). This suggests that the Y-linked genes affect sexually dimorphic phenotypes beyond the body size^25^, and may offer a resolution to sexual conflict more broadly in *C. maculatus*.

Roughly 50% of the Y genes have either exclusively X gametologs (99 genes, 58.6 Mb), or autosomal paralogs (157 genes, 1,153 Mb) with >80% protein sequence similarity (Fig. 2), indicating different origins for these genes. Notably the number of identified X and autosomal homologs is positively correlated (Pearson correlation: t = 20.55, df 113, p < 0.001) for 115 Y-linked transcripts that have homologs on both regions (Fig. S16, note that this includes also transcripts that additionally have homologs on uncharacterized contigs), suggesting their coupling to transposable elements. The higher absolute number of genes acquired from the autosomes follows the pattern seen in *D. melanogaster*^54^ and humans^14^, where all, or a substantial proportion of functional genes originate from autosomes, respectively. In contrast to known XY systems, in female heterogametic ZW taxa the W chromosomes mainly harbor Z gametologs^e.g.55^. The difference is explained by sexual selection favoring transpositions to the Y in male heterogametic taxa^14, 16, 54^. The finding that *C. maculatus* Y contains a mix of male-specific but also seemingly functional X gametologs, with high sequence similarity retained to the X, suggests that the ancestral X genes have still important roles in males and may evolve under purifying selection. These X gametologs are enriched for functions related to development and metabolic processes as well as RNA processing/splicing (Fig. S8). Sex-specific RNA splicing allows expression of alternative transcripts in the sexes and can facilitate sexual dimorphism^56, 57^. However, whether the X gametologs are dosage-sensitive and sexually concordant or sexually antagonistic and on the Y specialized for male beneficial functions remain to be tested.

The male-specific Y genes acquired via transposition from autosomes are functionally enriched for a broader range of terms than the gametologs, and account much of the general enrichment patterns observed across all Y genes (Fig. S8). An important avenue by which the Y chromosome can affect male phenotypes is by modulating gene expression throughout the genome, an effect described in *D. melanogaster*^58^ and for the *SRY* locus in mammals^59, 60^. In line with such a mechanism, Y-autosome epistatic effects have also been associated with sexually antagonistic coloration in guppies^61^. *C. maculatus* Y shows significant enrichment for DNA binding molecular function, a category that *SRY* also falls into (Fig. S7), which is consistent with the idea that the Y chromosome has a wider regulatory role. The autosomal paralogs on the Y are enriched for genes involved in protein translation & post-translational modification. While transcriptional sex differences are the commonly evoked explanation for how sexual conflict can be resolved^62^, this suggests that the Y chromosome may allow for translational modification to alter male beneficial phenotypic expression. The Y contigs also contain a large number of sequences with strong homology to other Y loci, suggesting frequent gene duplication events, which is commonly observed in Y chromosomes^14, 15, 17, 18, 20, 50, 63, 64^. The level of gene amplification we see is similar to the stickleback Y^20^, but less pronounced than in well characterized mammalian Y chromosomes^14, 15, 17, 18, 50, 64^. Amplification of genes on the Y may be fuelled by sexual conflict over associated traits^17^. Conservation of ampliconic genes is also observed in mammals^64^ and suggests that male specific amplified genes may have a large evolutionary advantage that withstand degenerative processes, such as Muller’s ratchet^64^. Ampliconic genes tend to be expressed in the testis and enriched for male-specific reproductive functions in mammals^14, 15, 17, 50^ and in stickleback^20^. But here we could not yet detect any genes with obvious reproductive functions in males, based on GO terms or previously described *C. maculatus* seminal fluid proteins (N=185)^65^.

X and Y do not form a chiasmata in *C. maculatus* and hence lack recombination by crossing-over and the PAR altogether. All species belonging to the *Callosobruchus* genus lack PAR based on their meiotic karyotypes^27^, suggesting that XY divergence predates the genus. Long evolutionary history without recombination with the X could explain why our analysis of sequence similarity between X and Y does not indicate synteny between them (Fig. S10), nor do we find indications for inversion blocks that could have led to recombination suppression between X and Y. Ampliconic regions are prone to structural rearrangements^14, 66^, which may cause further amplification and can also contribute to the lack of synteny between the X and Y.

### Molecular differences between the Y_S_ and Y_L_ haplotypes associated with body size variation

We detected substantial molecular differences between the sequences of the YS and Y_L_ haplotypes associated with two distinct male limited body size morphs^25^. As expected for a non-recombining, hemizygous genetic region, nearly all identified point mutation differences are fixed between the two Y haplotypes. The few detected SNPs could indicate real segregating polymorphisms (as the sample for sequencing consisted of multiple males) but are more likely artefacts caused by repetitive elements that add difficulty in genome assembly, accurate mapping, and variant calling. The 137 genes with fixed differences between the Y_S_ and Y_L_ haplotypes either in their coding or potential cis-regulatory regions are candidates to explain the phenotypic differences. Independent of body size, males with the Y_S_ haplotype also sire more offspring (*manuscript in preparation*). This could suggest that the Y haplotypes could cause differences in regulatory pathways associated with seminal fluid production or spermatogenesis, although we did not find any previously described *C. maculatus* seminal fluid proteins to be Y-linked. We identified *Drosophila* orthologs for 14 genes with fixed SNVs, and while we do not find any obvious causal candidate to explain the difference in reproductive capacity we find two notable orthologs: *male-specific lethal* 2 & *ubiquitin specific protease 47*. *Male-specific lethal* 2 is a well-known regulator of dosage compensation between the X and Y in *Drosophila melanogaster*^67^, which opens for differences in regulation of X-linked gene expression as a possible avenue to contribute to the phenotypic differences between the two Y haplotypes. Further, *ubiquitin specific protease 47* is known to interact with insulin/insulin-like signalling pathway (IIS) in *Drosophila*^68^, an important pathway that connects nutrient levels to metabolism, growth, development and longevity.

Remarkably, the longest *C. maculatus* Y chromosome contig also contains a *TOR* gene ortholog, a strong candidate to explain the Y linked size variation between the sexes as well as in males. The TOR signalling pathway is highly conserved and present in organisms from bacteria and plants to animals, and is one of the most ancient nutrient-sensing pathways^41^. The TOR pathway regulates growth and lifespan by coupling the growth factor signaling with nutrient sensing^41^. It is centrally involved in controlling cell metabolism, growth, proliferation and apoptosis. Together with IIS, TOR has also been implicated in differential gene expression between the sexes^69^ and more specifically sexual size dimorphism in *D. melanogaster*^57^. To our knowledge it has never been detected on a sex chromosome before. But the potential for Y specific regulation of the TOR pathway has recently been implicated in the male polymorphic Poecilid fish *P. parae*, where an inhibitor of the TOR pathway has been detected segregating on the Y^21^. To understand whether the *C. maculatus TOR* gene on the Y represents ancestral homology with the X, or has occurred by a transposition from the autosomes after X-Y divergence, we searched the genome for any *TOR* copies. We could not detect the *TOR* on the X, but only in one of the autosomal contigs, supporting its origin on the Y by translocation. We detected multiple transposable element sequences flanking the *TOR* sequence in all of the Y copies as well as the autosomal one (Fig. S15), which could have played a role in the translocation and should be subject to further investigations.

Importantly, we detected *TOR* copy number variation between the Y_S_ and Y_L_ haplotypes, which makes this gene the most likely candidate for the striking difference in male body size between the two haplotypes, and thereby sexual size dimorphism^25^. The elevated intronic sequence divergence between the autosomal and Y-linked *TOR* copies suggests that the translocation of the *TOR* has happened before the split of the two Y haplotypes, and also that sufficient time has passed since the translocation, to accumulate such differences in the introns. The exonic sequences have diverged only little, suggesting purifying selection on all copies to retain Y *TOR* regions as functional.

How the Y-linked *TOR* functions and putatively interacts with the autosomal *TOR* pathway presents a novel and exciting area for future research. The Y-linked, exonic *TOR* sequence is nearly identical to its autosomal paralog apart from five missing exons on the 5’end. The N-terminal of TOR proteins consist of HEAT repeats^70^ that mediate protein-protein interactions^71^. Empirical studies in *D. melanogaster*^72^ and amoeba *Dictyostelium discoideum*^73^ have demonstrated that overexpressing TOR inhibits cell growth and proliferation similar to loss of function mutants^72, 73^. Even a truncated extra copy of TOR is enough to reduce growth^72, 74^. Gene copy number can correlate positively with gene expression^75^, and *TOR* expression in adults in our data is overall male-biased (Fig. 3). Paralog interference has been suggested as one possible consequence of gene duplications^76^, whereby the paralogs can mutually exclude each other from binding with potential partners. The Y TOR could therefore affect male growth by interfering with the autosomal TOR pathway. The more common Y_S_ haplotype that makes males roughly 30% smaller than the Y_L_ haplotype has two additional copies of *TOR* on the Y, suggesting that the additional copies lead to further growth inhibition. Future work can establish how each of the copies in the two haplotypes may be expressed and function in regulating growth.

## Conclusions

Recombination is a key mechanism generating and maintaining allelic diversity across loci. Finding substantial diversity in the absence of recombination is therefore unexpected, but echoes a similar recent finding in a Y-polymorphic fish^21, 37^. Genetic variation should be rapidly fixed by selection and drift on the Y chromosome. Finding segregating Y polymorphism in both copy number and SNV differences in proximity to over 100 genes therefore suggests that processes such as frequency dependent selection likely have actively maintained these Y haplotypes in the population for a longer evolutionary time. Our identification of over 400 genes on the *C. maculatus* Y, despite the evidence of its degeneration, suggests that males can harbor substantial evolutionary potential through their Y chromosomes. We find that males with different body size morphs vary in the number of copies of a conserved growth factor gene that our data suggests has translocated from an autosome. The Y chromosome thus enables decoupling of the genetic response in the body size evolution between the sexes, positing the *TOR* pathway as the central regulator of sexual size dimorphism in *C. maculatus*.

## Methods

### Study organism and generation of the Y-lines

As a model organism to study sexual conflict, much is known about sexual antagonism in the seed beetle *Callosobruchus maculatus*. Aphagous adult *C. maculatus* females oviposit directly onto legume seed pods, within which larvae will develop for about 3 weeks, which allows for large scale experiments across multiple generations. The populations used in this study all stem from originally field caught (2010) individuals from Lomé, Togo (more details in^77^) and have been kept in the lab for ∼200 generations as 41 isofemale lines.

The creation of Y_S_ and Y_L_ haplotype introgression lines is described in detail in^25^. Briefly, replicated bi-directional selection was applied on male body size for 10 generations, giving rise to large (L) and small (S) males. We then crossed S and L males from these selection lines with females from a single inbred line (originating from the same Lomé base population as the selection lines, inbred for >20 generations^78^), respectively. At each subsequent generation, sons were backcrossed to females from the maternal line, for a total of 11 generations, after which the lines were sequenced. The backcrossing scheme replaced the original autosomes and the X chromosome with those from the inbred line while maintaining the non-recombining parts of the Y chromosome of the founding S or L males. Subsequent analysis of the body sizes confirmed the presence of two distinct male body size classes, while there was no difference in female body size^25^. We chose a single line representing each of the Y_S_ and Y_L_ haplotypes for sequencing.

### Sequencing and genome assembly

#### DNA extraction and library preparation

For the extraction of high molecular weight (HMW) DNA, we flash froze adult virgin males (within 24 h after emergence) in liquid nitrogen. Individual abdomens were dissected on ice to avoid thawing of the tissue by removing head, thorax and the elytra. 5 male abdomens were pooled and ground into fine powder with liquid nitrogen and a precooled pestle. The QIAGEN Genomic-Tip 20/g kit was used to extract HMW DNA, following the manufacturer’s protocol, with an over-night incubation time for cell lysis (i.e. 12 h). To achieve the required amounts of HMW DNA, we pooled 2 samples (total of 10 males per Y introgression line). 5 µg of genomic DNA were sheared on a Megaruptor3 instrument (Diagenode, Seraing, Belgium) to a fragment size of about 13-16 kb. The SMRTbell library was prepared according to Pacbio’s Procedure & Checklist – Preparing HiFi Libraries from low DNA input using SMRTbell Express Template Prep Kit 2.0 (Pacific Biosciences, Menlo Park, CA, USA). The SMRTbells were sequenced on a Sequel IIe instrument, using the Sequel II sequencing plate 2.0, binding kit 2.2 on three Sequel® II SMRT® Cell 8M per introgression line, with a movie time of 30 hours and a pre-extension time of 2 hours.

The genomes of the Y_S_ and the Y_L_ introgression lines were assembled individually using hifiasm (v. 0.7-dirty-r25)^79^ with default settings, yielding the S & L genomes. Haplotypes in the resulting assemblies were subsequently separated with purge_dups (v. 1.2.5, default parameters)^80^ and genome assembly completeness was assessed via BUSCO^81^, using default parameters on the insecta_odb10 database. We then chose the S genome (containing the ancestrally more frequently occurring Y_S_ haplotype) to be the reference genome, that we subsequently soft-masked for repetitive content^82^ using RepeatMasker (v. 4.1.2)^83^ and fully annotated using the BRAKER (v. 2.1.6)^84^ /TSEBRA (v. 1.0.3)^85^ pipeline. Detailed annotation methods are provided in the supplementary information: genome annotation.

### Identification of Y and X contigs

#### Mapping of putative X and Y contigs

We used gmap (v. 2021-03-08)^86^ default setting to quantify the number of hits of putative X and Y contigs, previously identified in *C. maculatus*^29^, to our assembled contigs in both genomes. Mapping to contigs shorter than 100 kb were excluded.

#### Sex assignment through coverage (SATC)

To independently identify putative sex-linked contigs in each of our assemblies, we employed SATC^38^, that uses normalised coverage information across contigs to first identify XX and XY individuals from sets of male and female samples^38^. Informed by a t-test, SATC compares normalised coverage at each contig between XX and XY individuals to find contigs with significantly different coverage. At sex-linked chromosomes, specific XY:XX coverage ratios are expected for X-linked contigs (0.5:1) and Y-linked chromosomes (1:0)^38^. In practice, coverage can be highly variable resulting in deviations from the strict expectation. Here we used the SATC approach to identify any contigs that had significantly different coverage between XX and XY individuals. We collected Illumina short-read sequencing data from Sayadi et al. (2019; ENA accession numbers: ERR3053159, ERR3053160, ERR3053163, ERR3053164, ERR3053161, ERR3053162, ERR3053165, ERR3053166)^29^. To increase certainty, we limited ourselves to the analysis of contigs > 100 kb in length. Contigs that are shorter than this account for a total of 24’939’325 bp and make up only 2.00% of the reference genome (1’246’713’675 bp). For more details see supplementary information: sex assignment through coverage (SATC).

#### PCR

We confirmed male-specificity of the longest identified Y contig with a multiplexed PCR, using two primers pairs, one that is Y specific and amplifies a 297 bp on utg000322l_1 product and a primer pair that amplifies an autosomal product of 189 bp length on utg000177l_1 as a positive control. For more detail see supplementary information: Molecular sexing.

#### Gene expression analysis

To assess how the identified sex-linked genes may be expressed we used gene expression data from virgin adult males (n=29) and females (n=32)^87^. The expression data was collected from reproductive tissues of the abdomen of virgin individuals from different inbred lines that originate from the same Lomé base population as the Y lines. We used splice variant aware mapping of transcript via STAR (v. 2.7.2b) with default settings^88^. We then used picard (v. 2.23.4)^89^ to mark duplicates and subread (v. 2.0.0) featurecount^90^ to summarize exons by gene IDs allowing for multimappers due to high gene duplications on the Y. Additionally, we also summarized exons by gene IDs using default setting (i.e. no multimapping). We then used DESeq2^91^ to analyse the gene expression patterns in males and females. We split the dataset into genes on the autosomes, and the identified X and the Y contigs.

#### GO enrichment on the Y vs the X

We examined how the Y chromosome is functionally diverged from the X using gene ontology enrichment analysis. For this we used the topGO R package^92^, with nodeSize parameter of 10, comparing the frequency of terms among the Y linked transcripts to those among all transcripts on the sex chromosomes (X and Y). Visualisation and clustering of the gene ontology enrichment analysis was done using ViSEAGO R package^93^, clustering GO terms based on Wang’s semantic similarity distance and ward.D2. Further aggregating of semantic similarity GO clusters was done with best-match average (BMA) method, as implemented in the ViSEAGO package.

#### Identification of Y homologs on the X and the autosomes

To identify whether the genes on the Y represent X gametologs or paralogs translocated from the autosomes, we used blastp (v.2.12.0)^39^ to compare Y proteins against all proteins in our reference assembly, requiring a minimum Y protein coverage of >50% and at least >80% AA identity, excluding self matches. To test whether X contigs show elevated number of Y gametologs due to shared XY ancestry, we compared the number of Y transcripts for which we exclusively find gametologs on the X, to the number of Y transcripts with exclusively paralogs on the autosomes, while accounting for difference in length or number of transcripts between autosomes and the X.

#### Y amplicons

The mammalian Y chromosomes contain large ampliconic regions enriched with high-identity segmental duplications e.g.^14^. Given that there seem to be a lot of duplicated genes also on the *C. maculatus* Y, we used blastn (v. 2.12.0)^39^ to blast all concatenated CDS for each transcript on the Y contigs against each other with stringent requirements of >95% query coverage and >99.9% sequence identity and excluding self matches, to detect amplicon groups on the Y.

### Characterisation of the Y_S_ and Y_L_ haplotypes

#### Variant calling

We used minimap2 (v. 2.18-r1015)^94^ to align reads to the reference allowing for up to 20% sequence divergence (asm20), as was recommended for HiFi reads. Duplicates have been marked with picard (v. 2.23.4)^89^. We then used DeepVariant (v. 1.3.0, default settings)^95^ with model type PACBIO for HiFi reads to call variants and GLnexus (v. 1.4.1)^96^ to merge the variant calling files for both Y haplotype lines. We used vcftools^97^ to filter the VCF files to only get single nucleotide variants (SNV) with a minimum depth of 5 and an upper depth cut-off of 65. SNV were categorized into single nucleotide polymorphisms that are shared between both Y introgression lines (SNP) and single nucleotide differences that are fixed between the two Y-introgression lines (fixed SNV).

#### TOR candidate gene analysis

A copy of a conserved gene coding for *target of rapamycin* (TOR) was discovered in a putative Y contig identified in the study that sequenced *C. maculatus* genome for the first time^29^. TOR is well known for its role in regulating growth across taxa (reviewed in^41^) and is therefore a prime candidate to explain the male body size difference between the Y_S_ and Y_L_ haplotypes. We identified and curated TOR locations in the genome (see supplementary methods ‘genome annotation’ for full details). To compare the different TOR regions we used MAFFT (v. 7.407, with –ep 0 –genafpair parameters)^98^ to align concatenated *TOR* CDS of each identified region in both assembled genomes and create a guided gene tree. To assess whether there are differences in exonic and non-coding sequences divergence across the different *TOR* regions we used fully factorial pairwise alignment of exonic and non-coding sequences separately. For the non-coding alignment, we masked identified exonic regions with bedtools maskfasta and then aligned the non-coding sequences with masked exons via Mummer nucmer (v. 4.0.0 with –maxmatch –c 100 parameters). For the exonic alignment we used the same procedure but masking the non-coding regions instead. We visualized the exonic and non-coding alignments with the R package circlize^99^.

## Data availability

Data and code will be made available upon acceptance on Dryad/Github repository.

## Supporting information

Supplementary methods and results

## Acknowledgements

We thank Johanna Liljestrand-Rönn for help and advice with DNA extraction, James M. Howie for help to develop the molecular sexing protocol and Madeline Chase for computational advice and feedback on figures. We thank Martyna Zwoinska and Göran Arnqvist for valuable discussions. The authors would like to acknowledge support of the National Genomics Infrastructure (NGI) /Uppsala Genome Center and UPPMAX for providing assistance in massive parallel sequencing and computational infrastructure. Work performed at NGI / Uppsala Genome Center has been funded by RFI / VR and Science for Life Laboratory, Sweden. Computations were enabled by re-sources in projects SNIC 2022/5-83 & SNIC 2022/22-176 provided by the Swedish National Infra-structure for Computing (SNIC) at UPPMAX, partially funded by the Swedish Research Council through grant agreement no. 2018-05973. This work was funded by the grants from the Swedish Research Council (grant no. 2019-05038) and Carl Trygger Foundation (grant no. CTS-18:163) to E.I. and a research grant from Nilsson-Ehle Endowments (grant no 146700723) to P.K.

## Author contribution

The study idea and general design was conceived by E.I. SATC, Repeat analysis and identifying *Drosophila* orthologs of Y-linked genes (incl. associated tables & figures) was done by R.A.W.W. Molecular sexing protocol was designed by K.P and E.I and optimized by K.P. Genome was assembled by C.T.R. and annotated by D.S. All other laboratory work, data analyses and figures were done by P.K. with input from E.I. The manuscript was written by P.K and E.I. with contributions from R.A.W.W. and D.S.

## Declaration of no conflicting interests

The authors declare no competing interests.

