## Supplementary methods and results for "Y-linked copy number polymorphism of target of rapamycin (TOR) is associated with sexual size dimorphism in seed beetles"

### Supplementary Figures & Tables

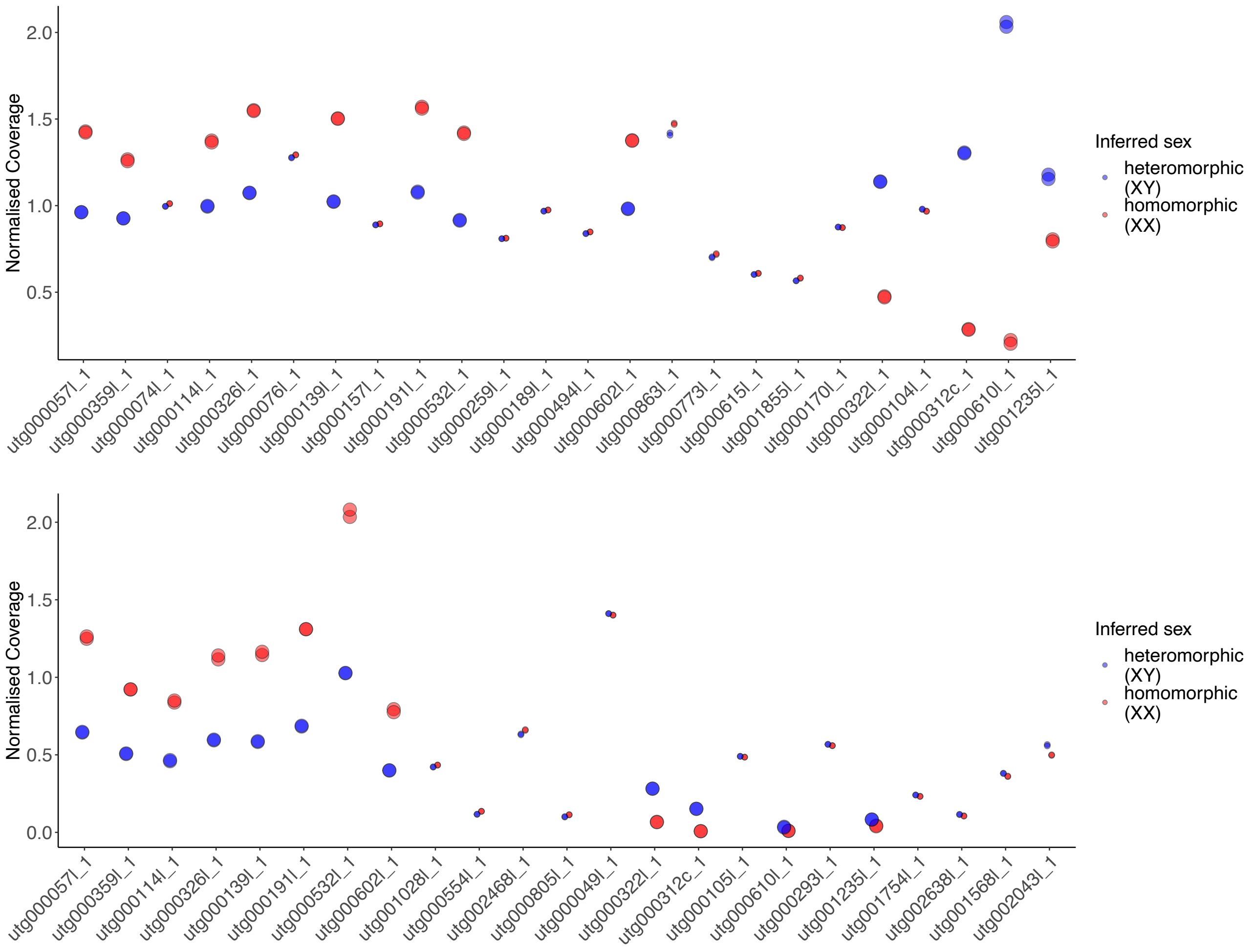

**Fig. S1| Sex assignment through coverage (SATC) for the S genome (containing the $Y_S$ haplotype).** Presented are all contigs that showed a significant coverage difference when mapping reads from two male (blue) or female (red) samples for un-masked (top) or repeat-masked $Y_S$ introgression line genome. Full details and stats in online Table O1.

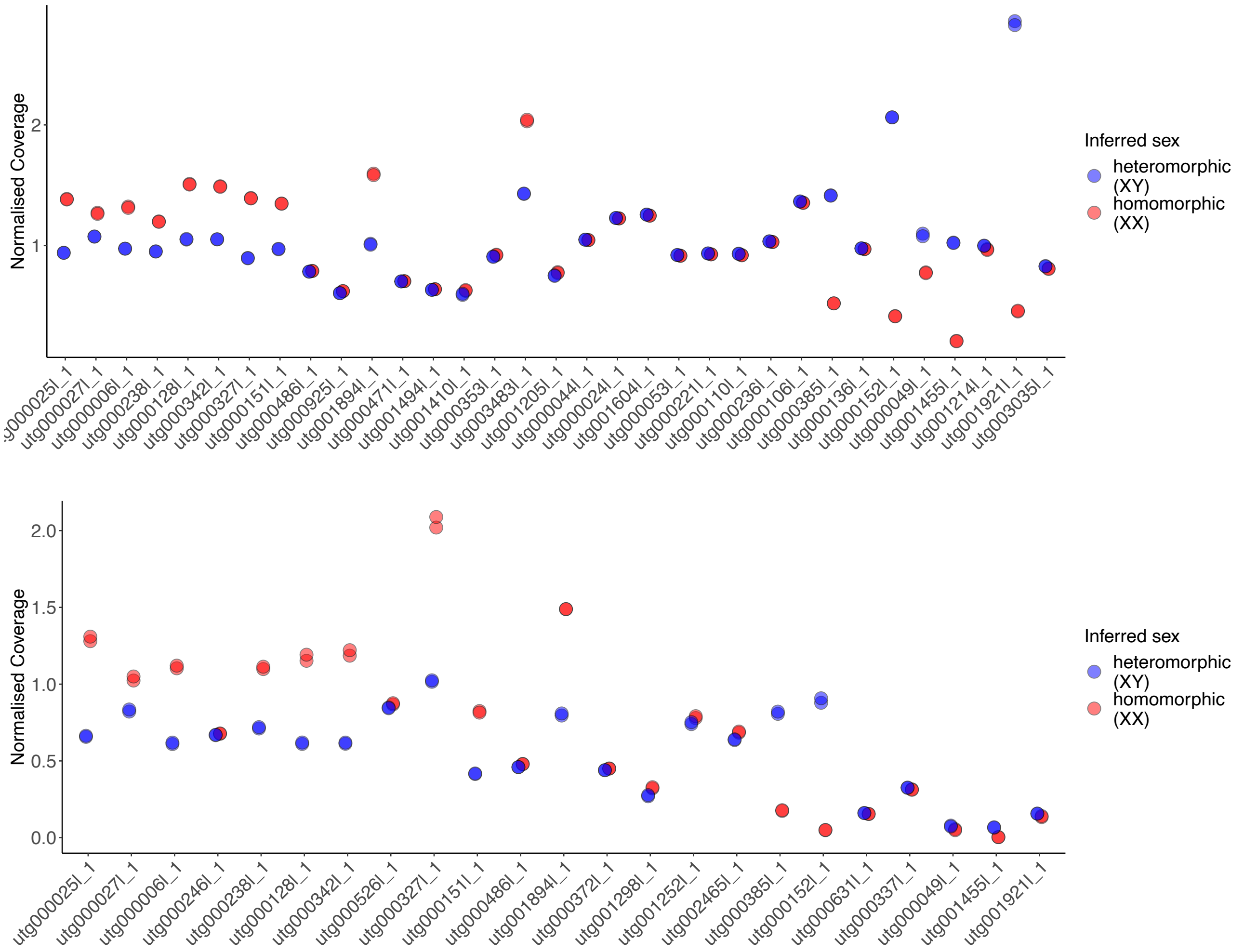

**Fig. S2| Sex assignment through coverage (SATC) for L genome (containing the $Y_L$ haplotype).** Presented are all contigs that showed a significant coverage difference when mapping reads from two male (blue) or female (red) samples for un-masked (top) or repeat-masked $Y_L$ introgression line genome. Full details and stats in online Table O1.

**Table S1| Overview of X and Y contig identification in the S genome.** Contigs indicated with checkmark show significant coverage difference of at least 10% for SATC and repeat masked SATC. Full details for SATC analysis are provided under Supplementary Methods & Results: Sex assignment through coverage (SATC). Contigs with checkmarks in gmap column showed high number of previously identified candidate X or Y reads mapping to the contigs. Note that PCR was designed to be utg003221_1 specific and does not apply for X contigs (see Fig. S3).

| contig | SATC | SATC repeatmasked | gmap | additional | Category | Length [bp] |
|---|---|---|---|---|---|---|
| *utg000322l_1* | ✓ | ✓ | ✓ | **PCR** | Y | 8432157 |
| *utg000312c_1* | ✓ | ✓ | ✓ | | Y | 800857 |
| *utg000610l_1* | ✓ | | | | Y | 667726 |
| *utg001235l_1* | ✓ | ✓ | | | Y | 211532 |
| *utg000057l_1* | ✓ | ✓ | ✓ | | X | 16805178 |
| *utg000114l_1* | ✓ | ✓ | ✓ | | X | 11442031 |
| *utg000139l_1* | ✓ | ✓ | ✓ | | X | 4942899 |
| *utg000191l_1* | ✓ | ✓ | ✓ | | X | 4268660 |
| *utg000326l_1* | ✓ | ✓ | ✓ | | X | 5268334 |
| *utg000359l_1* | ✓ | ✓ | ✓ | | X | 12192872 |
| *utg000532l_1* | ✓ | ✓ | ✓ | | X | 2462870 |
| *utg000602l_1* | ✓ | ✓ | | | X | 1174251 |

**Table S2| Overview of X and Y contig identification in the L genome.** Contigs indicated with checkmark show significant coverage difference of at least 10% for SATC and repeat masked SATC. Full details for SATC analysis are provided under Supplementary Methods & Results: Sex assignment through coverage (SATC). Contigs with checkmarks in gmap column showed high number of putatively identified X or Y reads mapping to the contigs. PCR was designed to be specific for $Y_S$ contig utg00322l_1 and does not apply for X contigs, note that utg00385l_1 and utg00152l_1 map to utg00322l_1 (see Fig. S14).

| contig | SATC | SATC repeatmasked | gmap | additional | Category | Length [bp] |
|---|---|---|---|---|---|---|
| *utg000049l_1* | ✓ | ✓ | | | Y | 280664 |
| *utg000385l_1* | ✓ | ✓ | ✓ | | Y | 2918857 |
| *utg001455l_1* | ✓ | ✓ | | | Y | 158603 |
| *utg001921l_1* | ✓ | ✓ | | | Y | 119672 |
| *utg000152l_1* | ✓ | ✓ | ✓ | | Y | 1415311 |
| *utg000006l_1* | ✓ | ✓ | ✓ | | X | 11724815 |
| *utg000025l_1* | ✓ | ✓ | ✓ | | X | 16843215 |
| *utg000027l_1* | ✓ | ✓ | ✓ | | X | 15046586 |
| *utg000128l_1* | ✓ | ✓ | ✓ | | X | 5276782 |
| *utg000151l_1* | ✓ | ✓ | | | X | 1168823 |
| *utg000238l_1* | ✓ | ✓ | ✓ | | X | 5546562 |
| *utg000327l_1* | ✓ | ✓ | ✓ | | X | 2601604 |
| *utg000342l_1* | ✓ | ✓ | ✓ | | X | 4997674 |
| *utg000486l_1* | ✓ | ✓ | | | X | 948900 |
| *utg001894l_1* | ✓ | ✓ | ✓ | | X | 897204 |

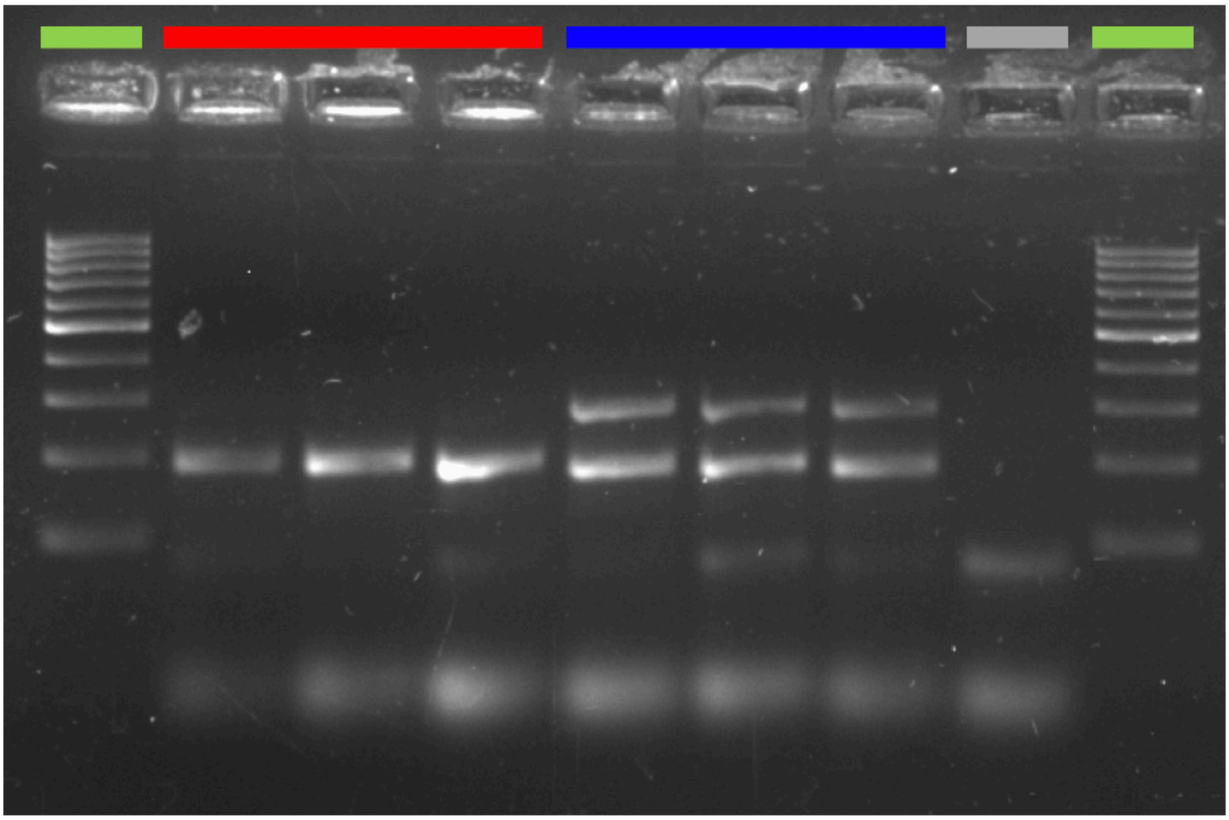

**Fig S3| Gel image of confirmatory multiplexed PCR amplification** of Y specific sequence (297 bp) in male (blue) but not in female (red) samples, while autosomal sequence (189 bp) amplifies in both sexes. The negative control (sample without DNA) shows no PCR amplification. 100 bp ladder (green) on both sides of the gel as a PCR product size reference. See detailed PCR protocol under Supplementary Methods & Results: PCR confirmation & molecular sexing protocol.

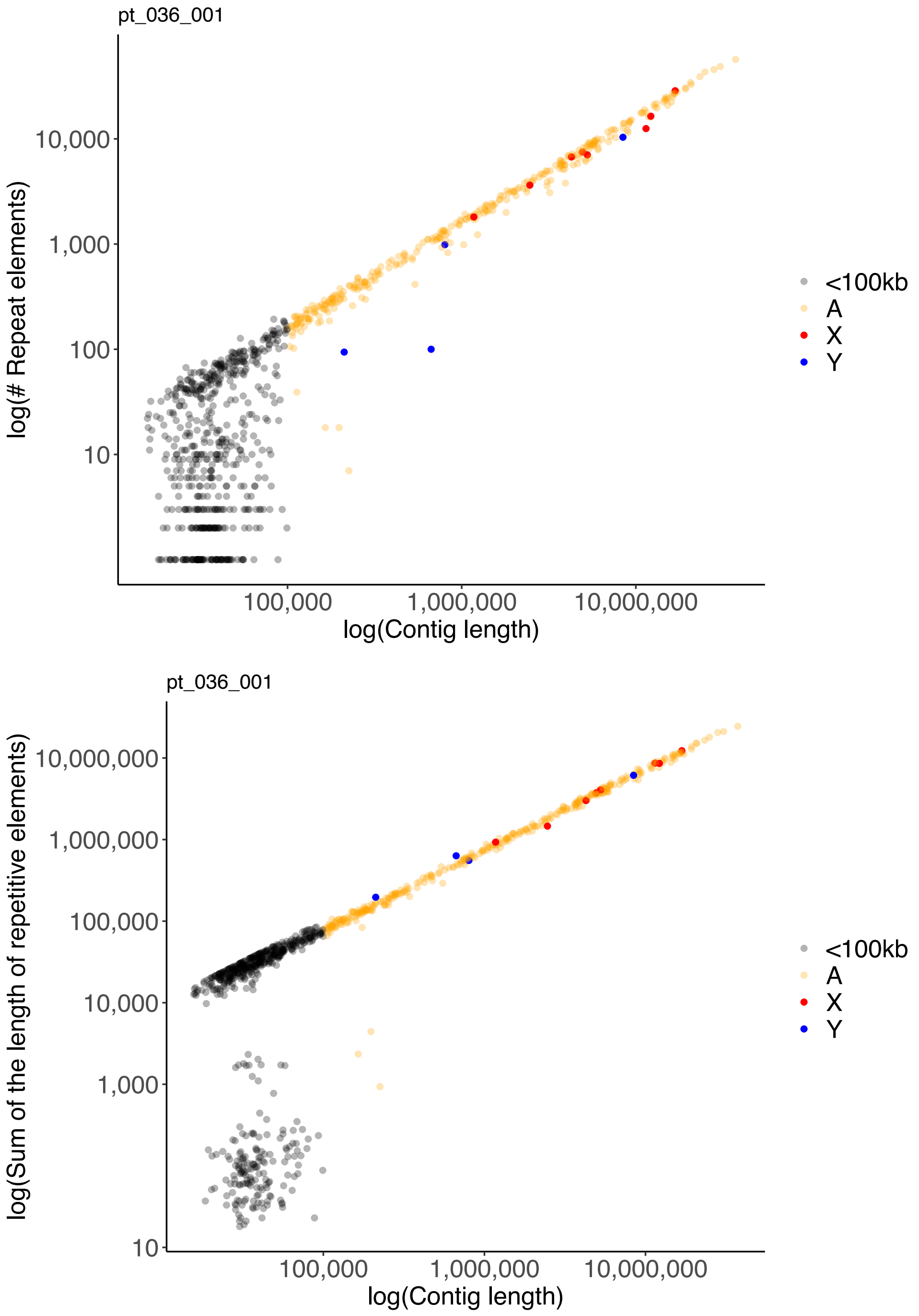

**Fig. S4| Repeat content in the Ys introgression line genome.** Identified repeat content per contig in number of repeats (top) or amount of bp covered by repeats (bottom). Contigs are color-coded by contig category: black = uncategorized contigs shorter than 100kb, yellow = autosomal contigs, red = X contigs, blue = Y contigs.

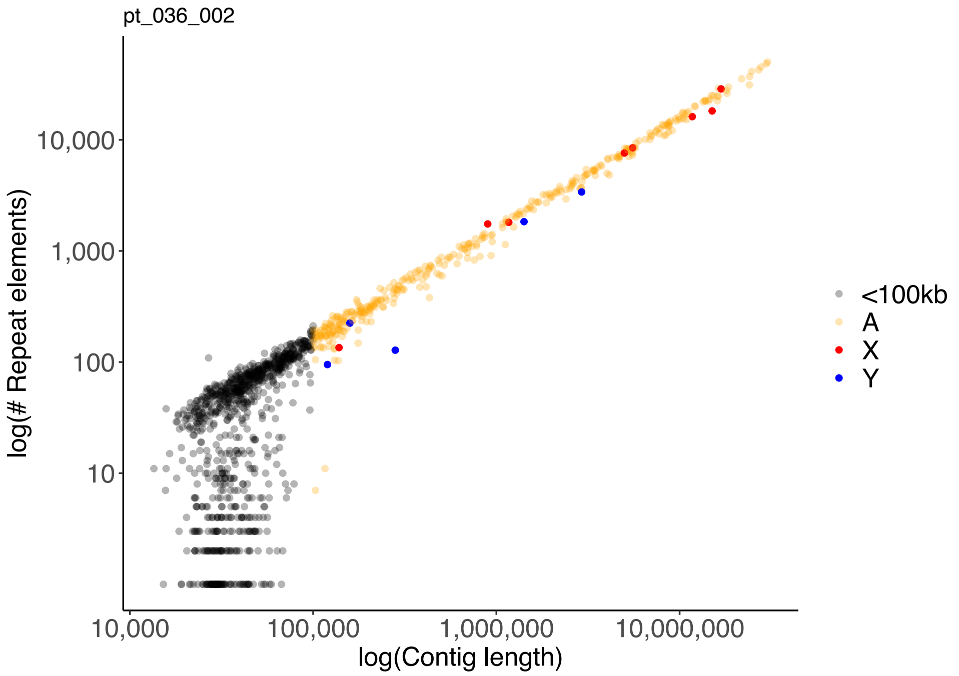

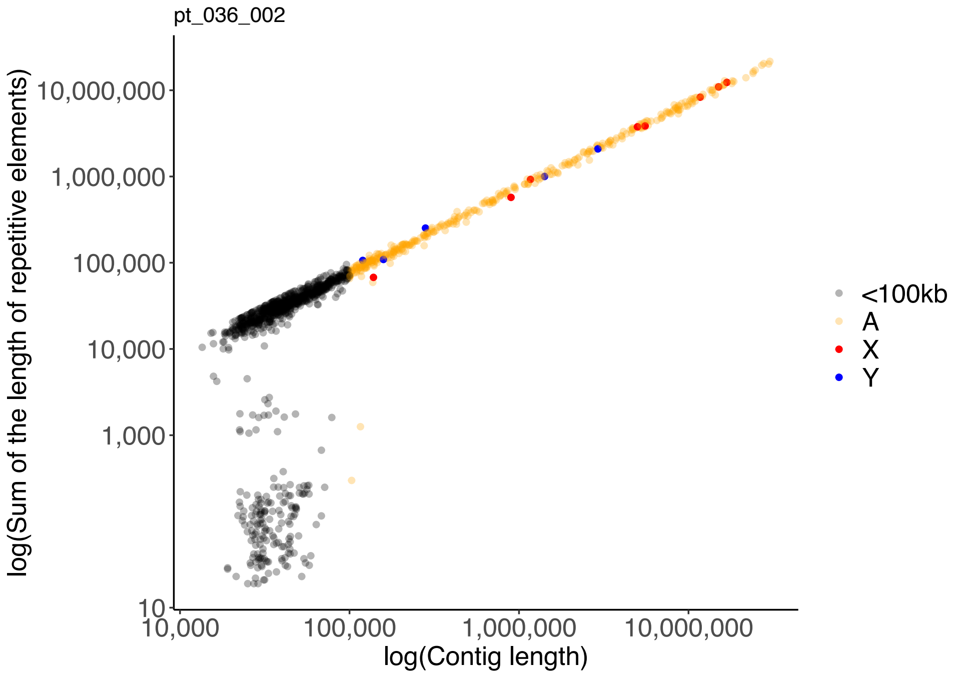

**Fig. S5| Repeat content $Y_L$ introgression line genome.** Identified repeat content per contig in number of repeats (top) or amount of bp covered by repeats (bottom). Contigs are color-coded by contig category: black = uncategorized contigs shorter than 100kb, yellow = autosomal contigs, red = X contigs, blue = Y contigs.

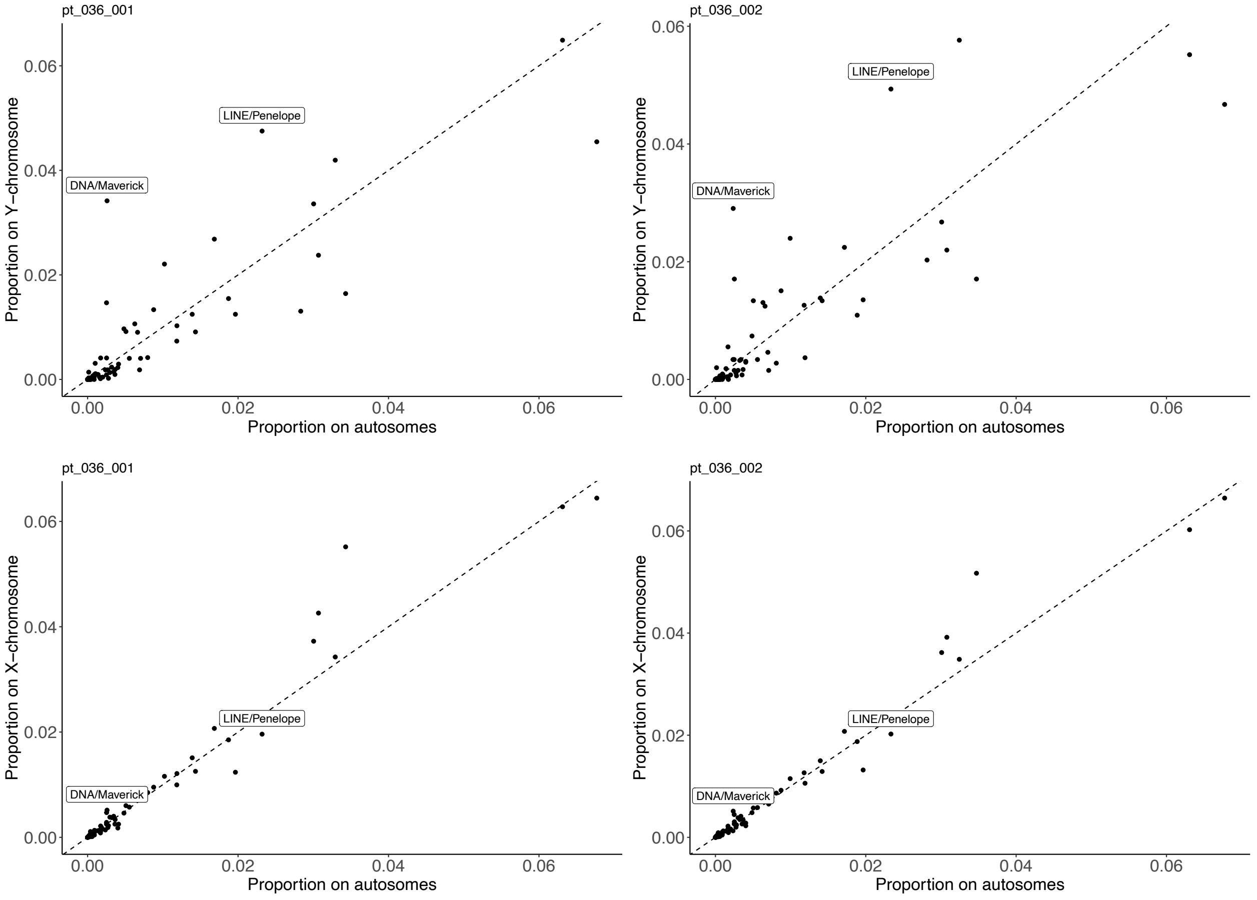

**Fig S6| Proportion of identified repeat categories** on sex chromosomes contigs (Y = top row; X = bottom row) compared to autosomal contigs for the S (left column) and L genome (right column).

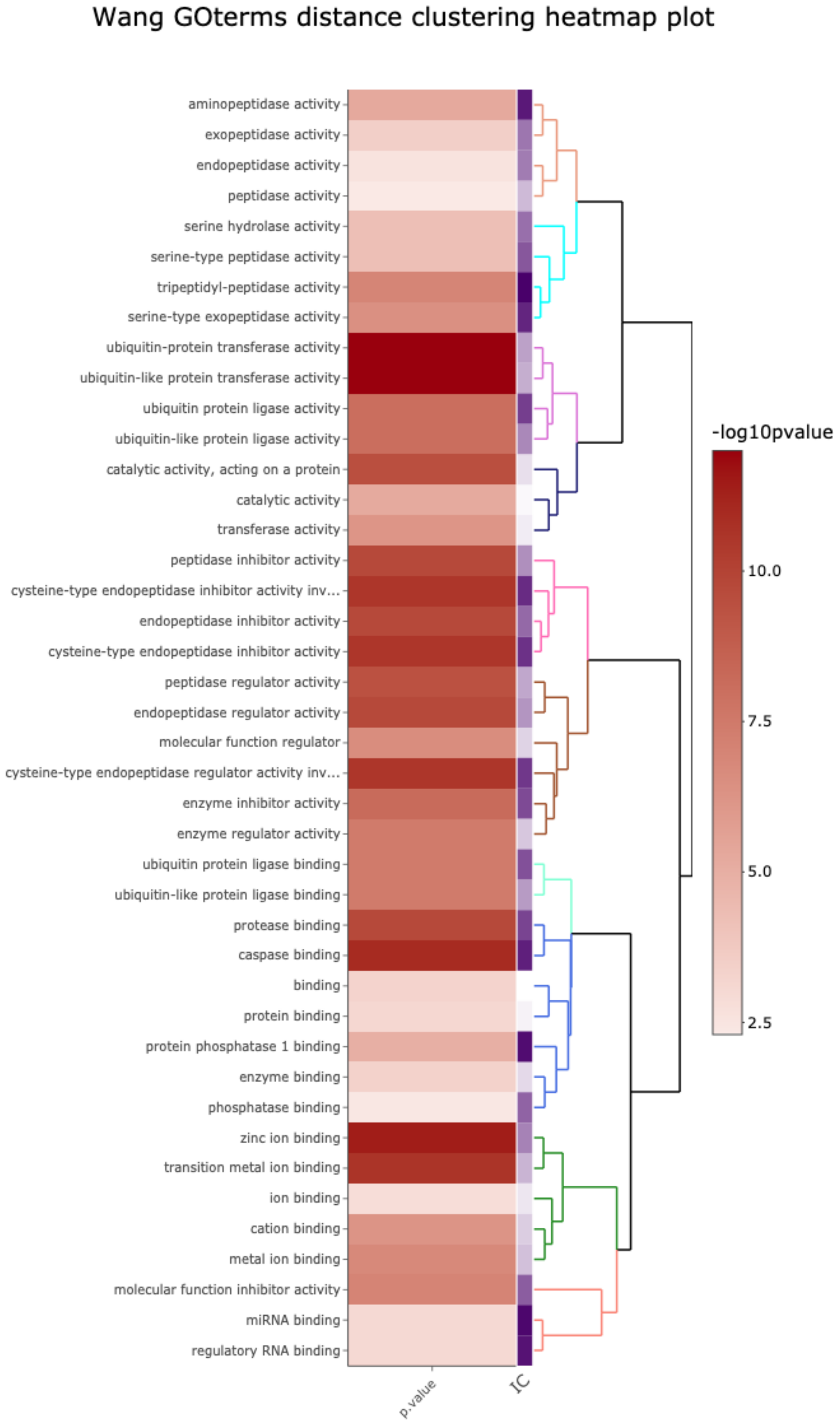

**Fig. S7| Gene ontology enrichment for $Y_S$-linked genes with 'molecular function' annotations.** Dendrogram of GO terms based on Wang's semantic similarity distance (nodesize = 8), heatmap (red) indicates statistical significance as -$\log_{10}$p-values (i.e. higher -$\log_{10}$p-values have higher statistical support) and information content (purple).

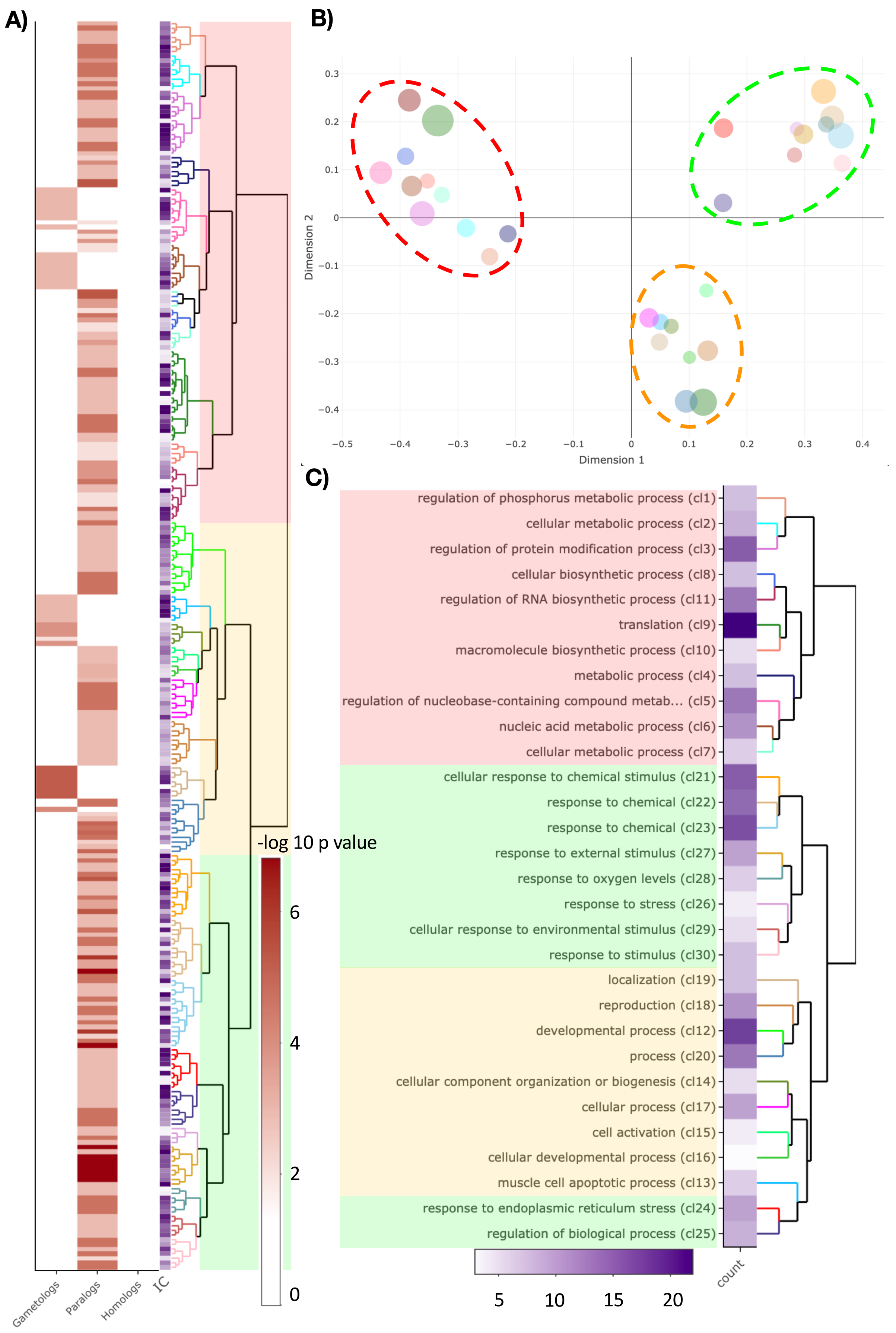

**Fig. S8| Gene ontology enrichment for $Y_S$-linked gene with X gametologs, autosomal paralogs, or homologs on both regions, compared to all Y-linked genes with 'biological process' annotations. A)** Dendrogram of GO terms based on Wang's semantic similarity distance, heatmap (red) indicates statistical significance as -$\log_{10}$p-values (i.e. higher -$\log_{10}$p-values have higher statistical support) split by gene homology (X homology = gametologs, autosomal homology = paralogs, both X and autosomal homology = homologs) and information content (purple), node size parameter = 5. **B)** Multi-Dimensional Scaling (MDS) plot based on Best-Match Average (BMA) distance, representing the proximities of dendrogram clusters in (**A**). Dot size indicates the number of GO terms within each cluster. We highlight the three major functional groups with dashed ellipsoids (green ≅ response to stimulus; yellow ≅ developmental processes; red ≅ metabolic processes) **C)** Dendrogram representation of clusters from (**B**) with GO term description of the first common GO ancestor and heatmap for the number of GO terms within each cluster.

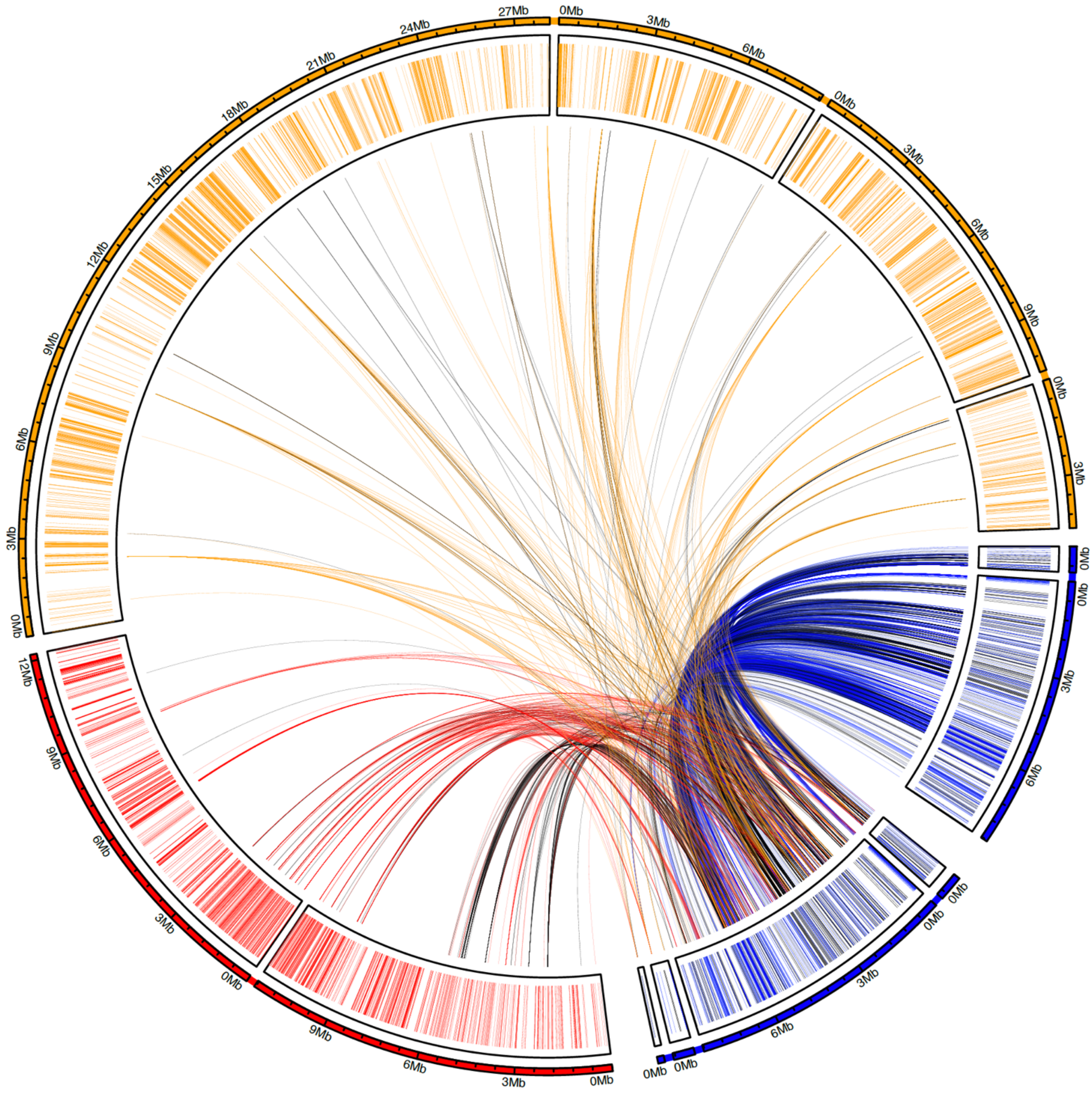

**Fig. S9| Gametologs, autosomal paralogs and Y duplications.** Overview of Y paralog distribution, showing the top 8 contigs with the most unique Y paralogs, each protein match to Y genes is shown with the genomic link. The highest number of unique Y homologous sequences are on the Y contigs (utg000322l_1 & utg000312l_1) themselves, suggesting a general pattern of gene duplication on the Y. Outer track shows the relative contig length (A, X and Y contigs are highlighted in yellow, red and blue, respectively), indicated are all gene positions in the respective color. Links and genes, shown in black indicate that matching Y gene is in a region that has been masked by RepeatMasker.

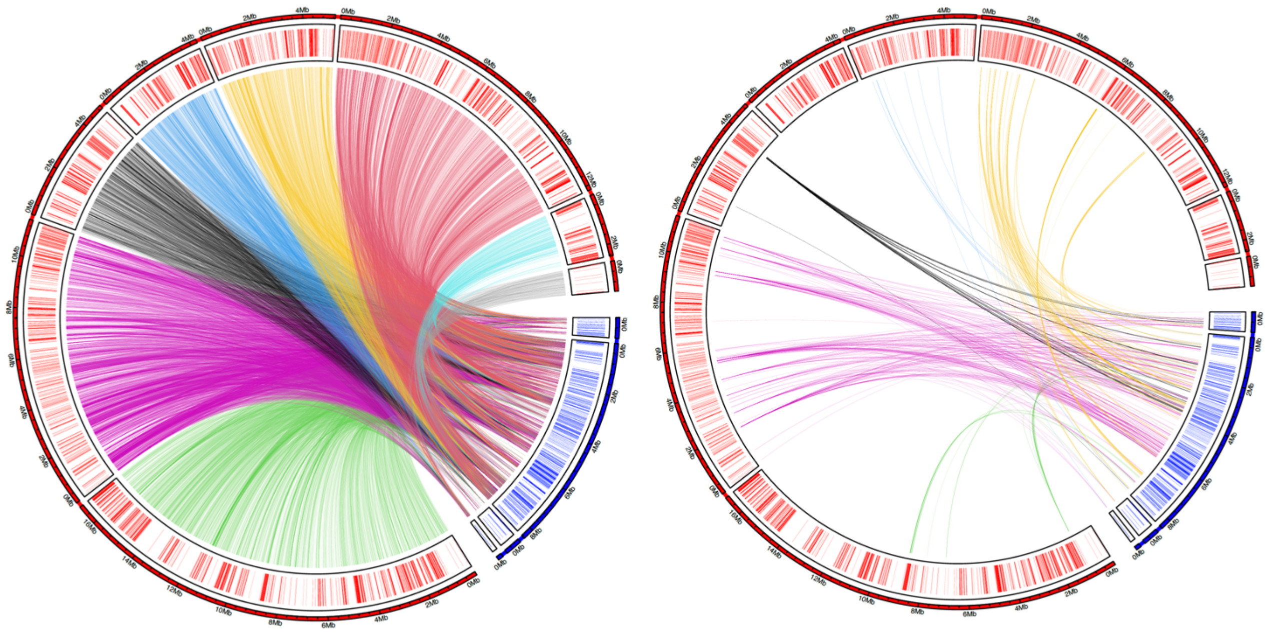

**Fig. S10| No apparent synteny blocks between X and Y contigs.** Genomic links between X (highlighted in red) and $Y_S$ (highlighted in blue) contigs represent nucleotide sequence similarity (left) and distribution of identified gametalogs (right) between the two sex chromosomes, gene positions are indicated with bars. Despite high nucleotide sequence similarity between X and Y there are no clear synteny blocks or inversions.

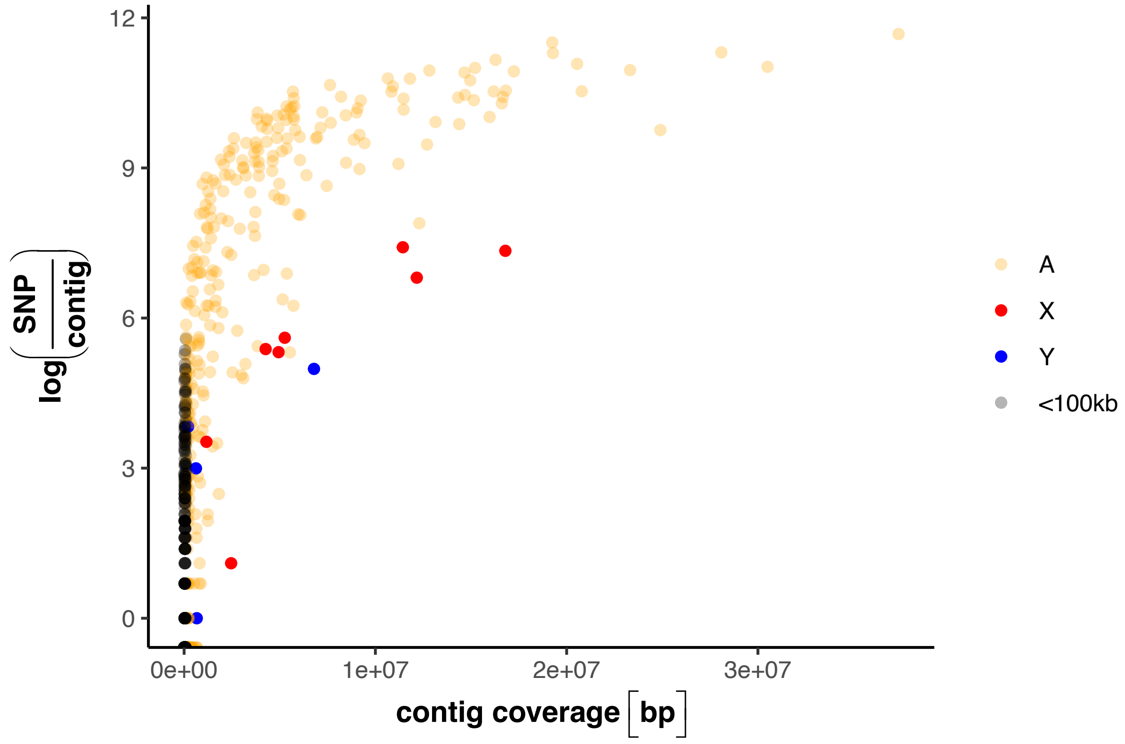

**Fig. S11| Number of SNPs per contig plotted against the length of the contig**, color coded by contig category (i.e. autosomal contigs in yellow, X contigs in red, Y contigs in blue and uncategorized contigs shorter than 100kb in black). Note that contig length considers only bp with coverage when mapping the $Y_L$ and $Y_S$ introgression line genomes against each other.

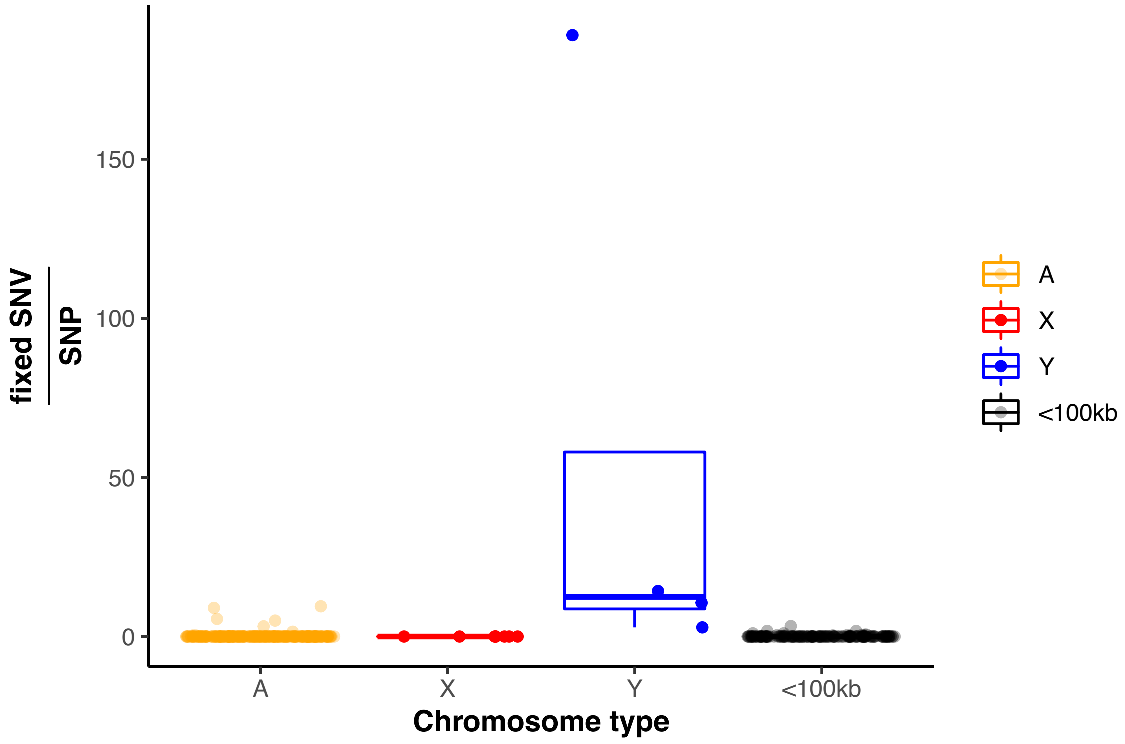

**Fig. S12| Ratio of fixed SNV over SNPs** per contig and contig category.

**Table S3| Identified Drosophila orthologs from FlyBase** for genes with fixed SNV between the two *C. maculatus* Y haplotypes.

| **FlyBase gene ID** | **gene name** | **gene symbol** |
|---|---|---|
| FBgn0034802 | CCHC-type zinc finger nucleic acid binding protein | CNBP |
| FBgn0262534 | - | CG43088 |
| FBgn0263400 | - | CG43446 |
| FBgn0031318 | - | CG4887 |
| FBgn0261647 | AXIN1 up-regulated 1 | Axud1 |
| FBgn0260635 | Death-associated inhibitor of apoptosis 1 | Diap1 |
| FBgn0020370 | Tripeptidyl-peptidase II | TppII |
| FBgn0020370 | Tripeptidyl-peptidase II | TppII |
| FBgn0005616 | male-specific lethal 2 | msl-2 |
| FBgn0016756 | Ubiquitin specific protease 47 | Usp47 |
| FBgn0034948 | Protein phosphatase 1 regulatory subunit 15 | PPP1R15 |
| FBgn0036668 | Zinc finger CCHC-type containing 7 | Zcchc7 |
| FBgn0037105 | Site-1 protease | S1P |
| FBgn0020370 | Tripeptidyl-peptidase II | TppII |

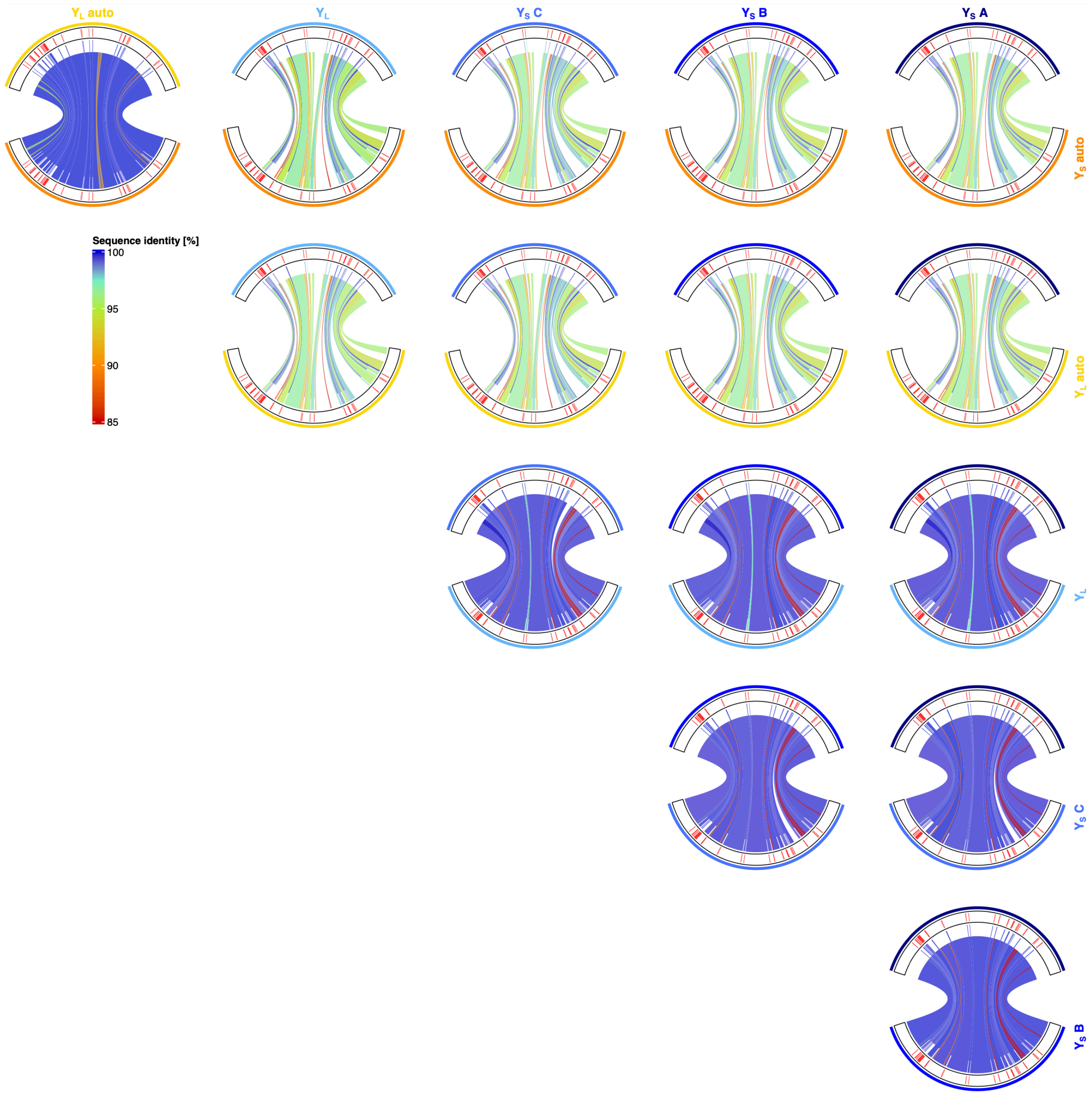

**Fig S13| Fully factorial pairwise TOR region comparison.** Nucleotide alignment with Mummer of coding (exonic) and non-coding regions separately. The outer track in red indicates the *TOR* exon positions. Genomic links show nucleotide similarity, exonic links are shifted upwards, while non-coding alignments are shifted down to ease distinction between the regions. Comparison of the autosomal (A) and Y-linked (Y) shows structural differences (gaps) and low sequence identity matches in the non-coding regions, while the exonic regions seem conserved between A & Y. Note that all Y *TOR* regions lack the exon 1-5 (from the 5' end) and exon 6 is only partially present. Exon 7-25 are present in all Y *TOR* regions. When comparing autosomal ($A_S$ vs $A_L$) or Y-linked ($Y_S$ vs $Y_L$) *TOR* regions across the two assembled genomes, we find high nucleotide similarity in both non-coding and exonic regions alike.

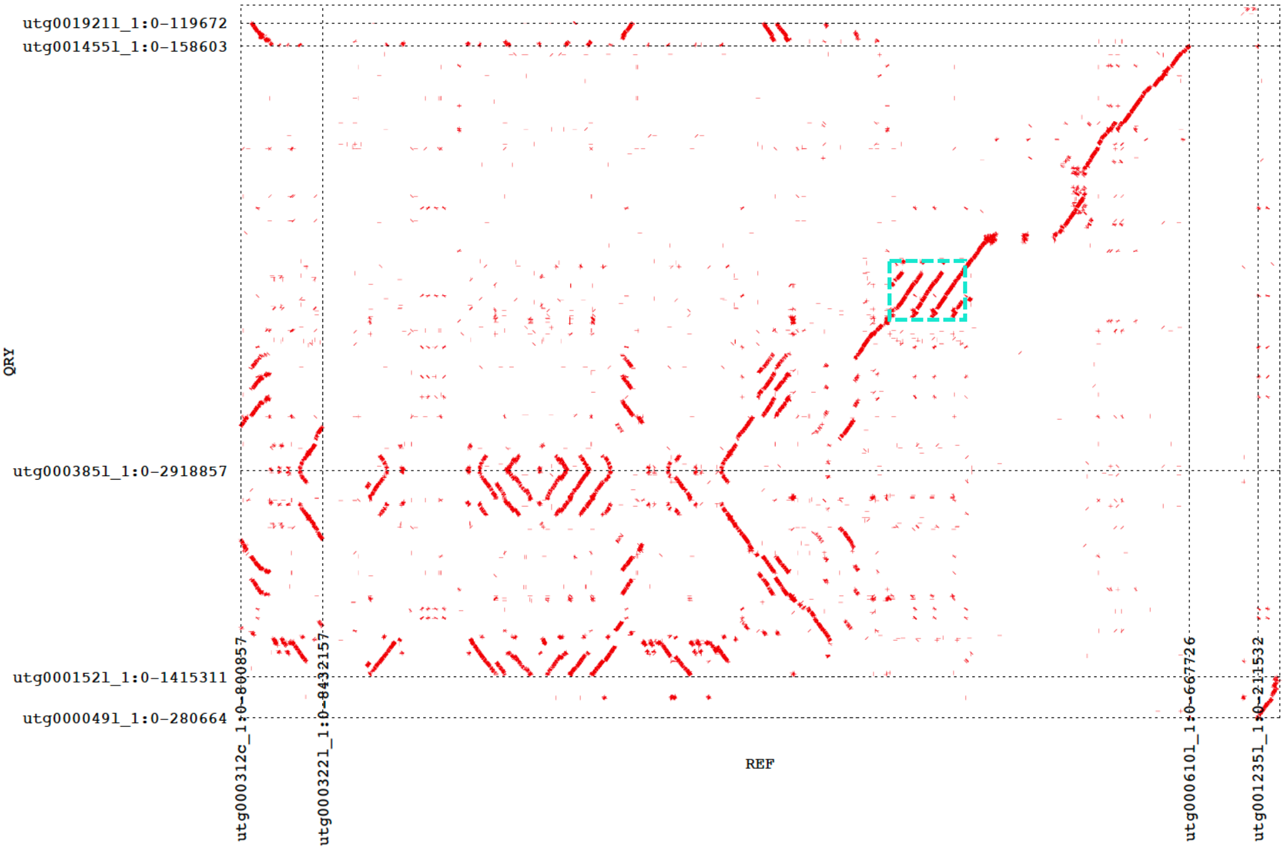

**Fig. S14| Y contig alignment between $Y_S$ (x-axis) and $Y_L$ (y-axis) haplotype.** The TOR region is highlighted in turquoise. Multiple diagonal lines (stacking on the x-axis) in the TOR region indicate that the TOR region exists in three copies on the $Y_S$ haplotype contig, used here as the reference, but only once on the $Y_L$ haplotype contig.

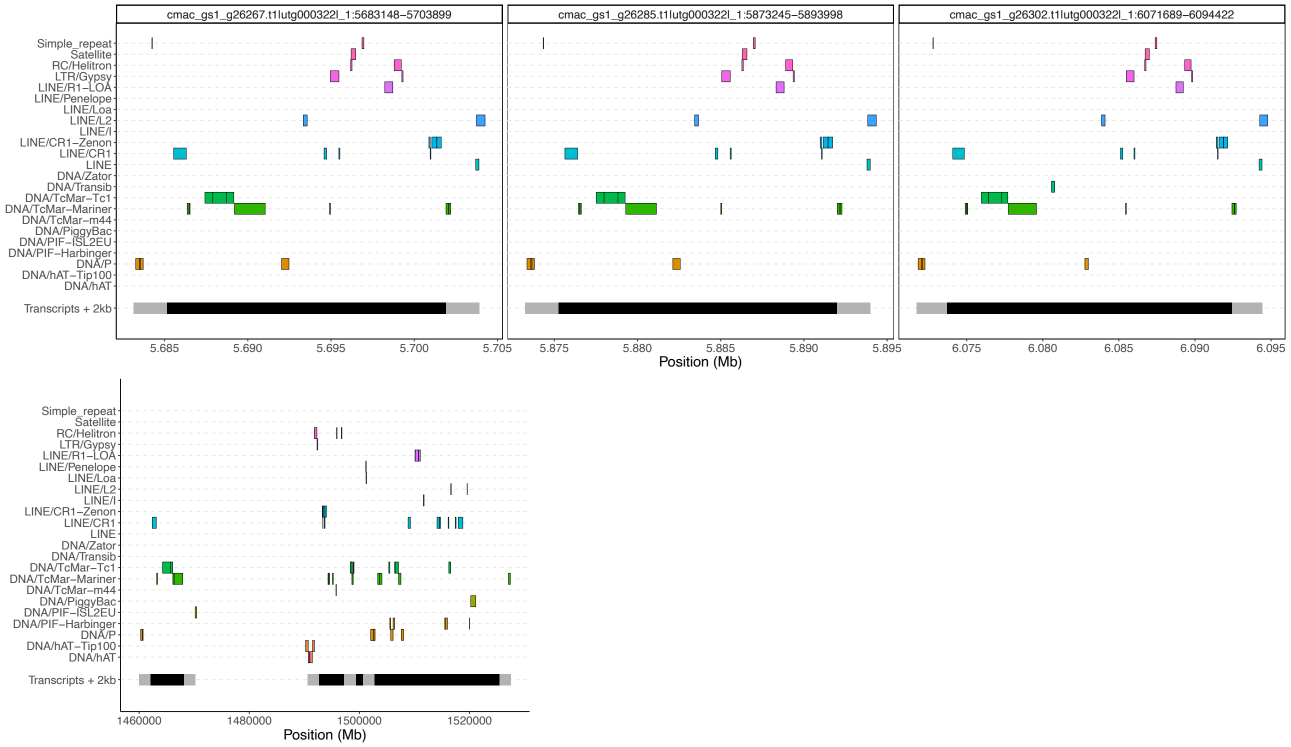

**Fig. S15| TOR regions in the $Y_S$ genome and identified repetitive elements within or in close proximity to the TOR region.** Distribution of repetitive elements in the Y-linked TOR regions (top) are very similar among each other and show some similarities with the autosomal TOR region (bottom).

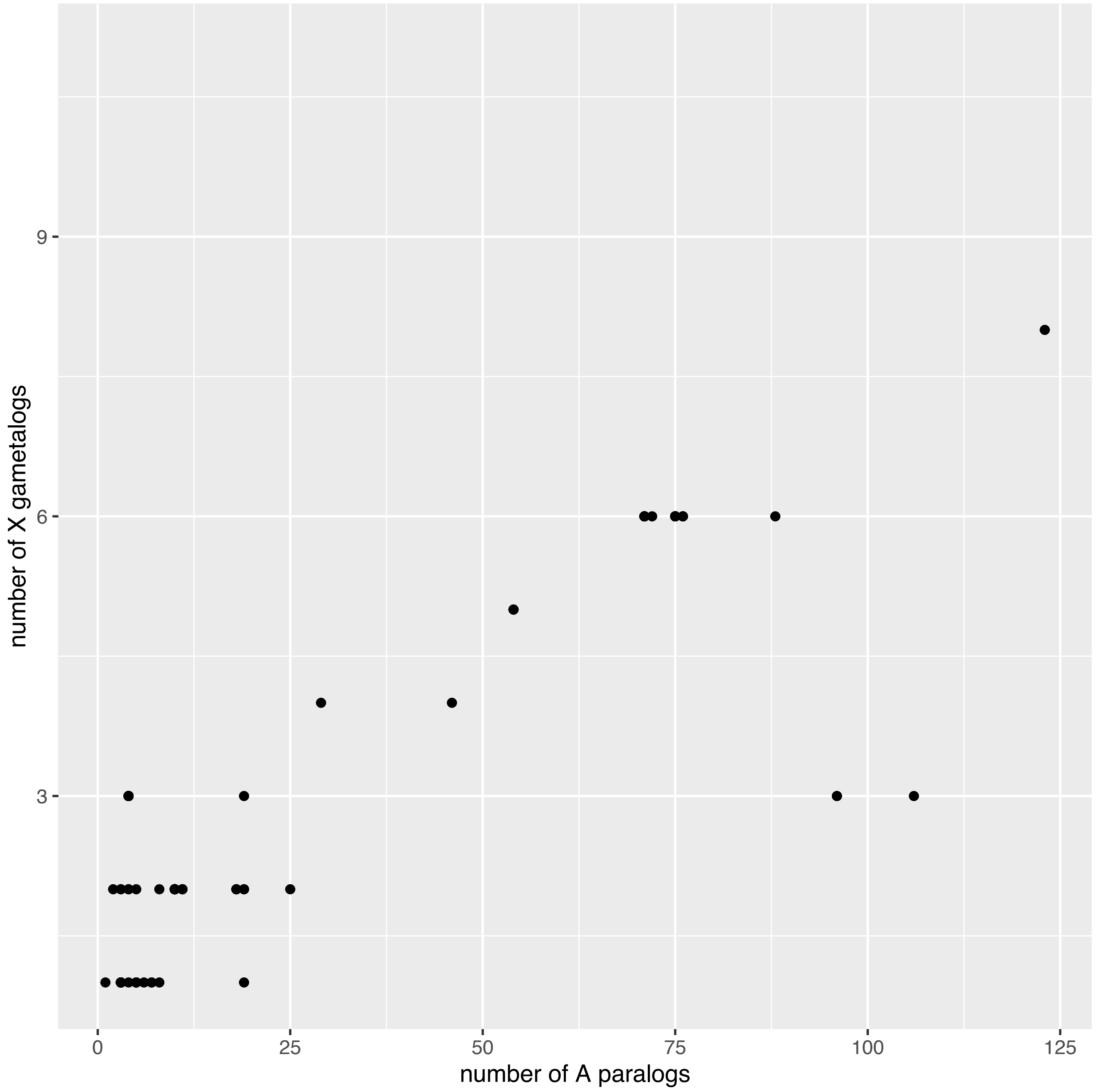

**Fig. S16| Y-linked transcripts with homologs on both X and autosomes** show a positive correlation. Y-linked transcripts with a high number of gametologs (Y axis) are more likely to also have a high number of autosomal paralogs (X axis).

### Supplementary Methods & Results:

#### Genome annotation

The primary-1 genome (i.e. $Y_S$ line males) was soft-masked for repetitive content using RepeatMasker 4.1.2-p1[1] in sensitive mode (options '-xsmall -s -u -engine ncbi -gff') with an existing curated repeat library for *C. maculatus* [2] & (A. Suh & J. Galbraith, pers. comm.). A GFF-format file containing the coordinates and identities of masked repeats will be made available later.

The soft-masked primary-1 genome was then used as the reference for gene model annotation with the BRAKER/TSEBRA pipeline, using BRAKER 2.1.6 [3] and TSEBRA 1.0.3 (through commit 336c380 dated 2021-11-26, https://github.com/Gaius-Augustus/TSEBRA; [4]). This pipeline uses separate steps for de novo determination of gene models in concert with RNA-Seq data (BRAKER1) and with protein alignments (BRAKER2), and a final step (TSEBRA) that synthesizes information from each to produce a final set of gene models.  Species-specific gene model training was performed with AUGUSTUS 3.4.0 [5].

Input RNA-seq data (**Paper IV**) were aligned to the reference genome using STAR 2.7.9 [6], with BAM output sorted by coordinate. The BRAKER1 step uses the RNA-Seq alignments to train de novo gene finding with GeneMark-ET and AUGUSTUS [5,7–13]. BRAKER was run with the options '--softmasking --UTR=off' and '--bam' followed by the list of BAM files containing aligned RNA-Seq reads.

Input protein data were:

- Proteins from the previous *C. maculatus* 1.0 genome assembly (ENA accession PRJEB30475, [2])
- Proteins from the genome releases of five beetle taxa belonging to the same family as *C. maculatus* and two closely related families, downloaded from I5K (https://i5k.nal.usda.gov/): *Leptinotarsa decemlineata*, Chrysomelidae (GCF_000500325.1); *Diabrotica vergifera*, Chrysomelidae (GCF_003013835.1); *Dendroctonus ponderosae*, Curculionidae (accession GCF_000355655.1); *Sitophilus oryzae*, Curculionidae (GCF_002938485.1); *Anoplophora glabripennis*, Cerambicidae (GCF_000390285.2).
- The complete Arthropoda section of OrthoDB 10v1 (https://www.orthodb.org/; [14]).

The BRAKER2 step uses genomic alignments of the protein sequences and ProtHint 2.6.0 plus de novo gene finding with GenMark-EP+ [15–18]. BRAKER was run with the options '--softmasking --UTR=off --epmode' and '--prot_seq' followed by the file containing the combined protein sequences.

The TSEBRA step combined gene models from the BRAKER1 and BRAKER2 steps, weighting the contributions of various sources of evidence to produce the final set of gene models. Following TSEBRA recommendations, the GTF gene identifiers from BRAKER1 and BRAKER2 were adjusted with TSEBRA's 'fix_gtf_ids.py' script. We used default weighting values when combining evidence.  We examined an alternate set of configuration values which favored high-quality protein alignments and reduced the contribution of 'good' protein alignments and of RNA-Seq data (configuration options 'P 0.05 E 6'), but this made no change to the number of gene models and very little change to transcript numbers.

One result of the TSEBRA step is that occasionally, nonoverlapping gene models will be annotated as alternate transcripts of the same gene.  A similar difficulty arises when a single gene in vitro is annotated as separate genes, or separate transcripts.  As a result, we consider our functional annotation unit to be the transcript (entries marked 'transcript' in GFF files) rather than the gene ('gene' entries).

Functional annotation of protein sequences extracted from the genome using the final gene models, including appropriate GO terms, was produced with eggNOG v5 and emapper 2.1.6 (http://eggnog-mapper.embl.de/; [19,20].

To produce targeted annotation for the serine/threonine kinase mTor-related gene models, we searched GenBank for candidate sequences using the human mTor isoform 1 protein (GenBank accession NP_004949.1, 2549 aa). We selected several candidate sequences from species within Coleoptera suborder Polyphaga, all ~2400 aa (GenBank accessions AKB11618.1, ALE20544.1, CAH0563318.1, CAH0563403.1, CAH1377105.1, KAF2880605.1, KAF5280820.1, KAF5285925.1, KAF7282696.1, RZC37432.1, UIB01653.1, XP_971819.1, XP_017768823.1, XP_018572076.1, XP_019880827.1, XP_022907797.1, XP_023015005.1, XP_025831250.1, XP_028145210.1, XP_030750054.1, XP_031352545.1, XP_044759281.1, XP_045478375.1). Hits from *C. maculatus* included two shorter sequences (VEN43112.1, 1786 aa; VEN51984.1, 1011 aa) not specifically annotated as mTor. We aligned all of these sequences with MUSCLE 3.8.1551[21] (options -diags -clwstrict). Visual inspection showed that the two *C. maculatus* sequences overlapped at their 3' and 5' ends, respectively, and together represented nearly the complete length of the mTor alignments. We separately aligned the two *C. maculatus* sequences with MUSCLE as above and used the **cons** tool from EMBOSS 6.6.0.0[22] (options -identity 1 -setcase 0.5) to produce a 2437-aa candidate mTor sequence for *C. maculatus*. The candidate was aligned to the repeat-masked genome using **exonerate**'s protein2genome model[23] (options --percent 80 --showtargetgff), which found four high-quality gene models, each with identity ≥99.67%. One complete gene model was found on an autosomal contig, and three gene models were found all within ~435 kbp on a $Y_S$ contig, all missing the first 620 residues of the candidate. We named the autosomal model mTor and the $Y_S$ contig models, in coordinate order, yTor-A, yTor-B and yTor-C. We adjusted the gene model descriptions to be consistent with the BRAKER/TSEBRA annotation and added functional annotations with eggNOG v5[19] and emapper 2.1.6[20] (http://eggnog-mapper.embl.de/). We incorporated these four gene models into the full annotation by removing overlapping gene models in the same orientation only; all mTor and yTor models were in '-' orientation and overlapped with other gene models in '+' orientation.

### Sex assignment through coverage (SATC)

**Methods:**

We collected available short-read sequencing data from 2 males and 2 female individuals from a previous study ([2]; ENA sample accession numbers: SAMEA5215851, SAMEA5215852, SAMEA5215853, SAMEA5215854, SAMEA5215855,SAMEA5215856, SAMEA5215857, SAMEA5215858). We concatenated reads from the same individuals into single fastq files for each individual and mapped these to our PacBio HiFi assembly using bwa mem (v. 0.7.17; [24]). We filtered the resulting .bam file with samtools (v. 1.14; [10]) to remove reads that were not properly paired, as well as all supplementary alignments of reads. The *C. maculatus* genome is extremely repeat-rich [2], and sex-chromosomes in general may be expected to be very repeat-rich [25]. We therefore also additionally filtered the data to avoid undue influence of high coverage repeat regions. We used RepeatMasker (v. 4.1.2; [1]) to first identify regions of repetitive DNA from the *C. maculatus* genome assembly using a curated *C. maculatus* repeat sequence library (A. Suh & J. Galbraith, pers. comm.) and then used bedtools (v. 2.29.2; [26]) to remove alignments to those regions from bam files. Finally, for both unfiltered and filtered .bam files we computed coverage statistics from the final bam files with samtools indxstats.

**Results:**

The analyses of filtered and unfiltered coverage data for both the $Y_S$ and the $Y_L$ genome, SATC correctly identifies the sexes of samples. Female samples (CmacF1 and CmacF2) are both inferred to be the homomorphic sex, while the two male samples (CmacM1 and CmacM2) are both inferred to be heteromorphic (Fig. SATC). SATC identified several contigs with significant coverage differences (Fig. S1-2), however, in some cases the effective coverage difference between the male and females samples, while statistically significant, was low (< 10% coverage difference) and we did not further consider such contigs on the basis of the SATC analysis alone (see online Table O1 for full SATC details). None of the identified contigs did pass a stringent Bonferroni correction for multiple testing, including contigs with additional, independent evidence (Table S1-2).

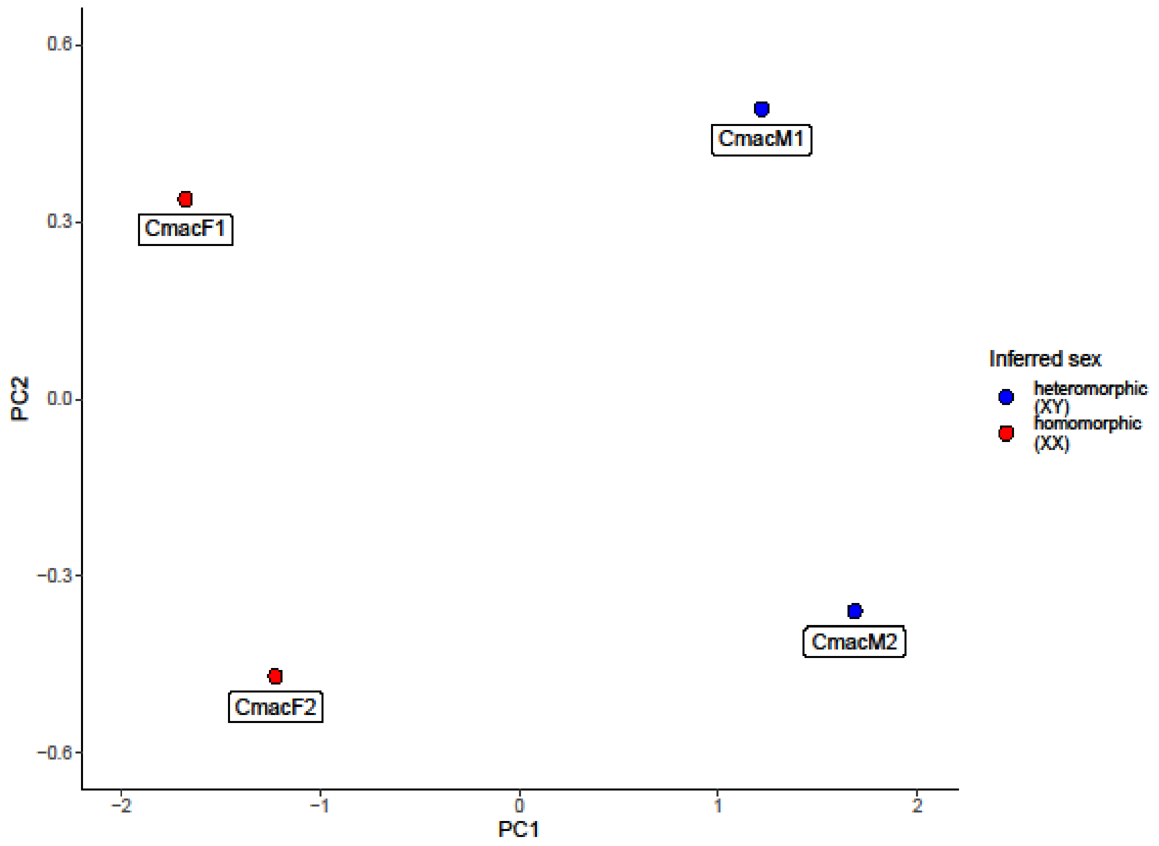

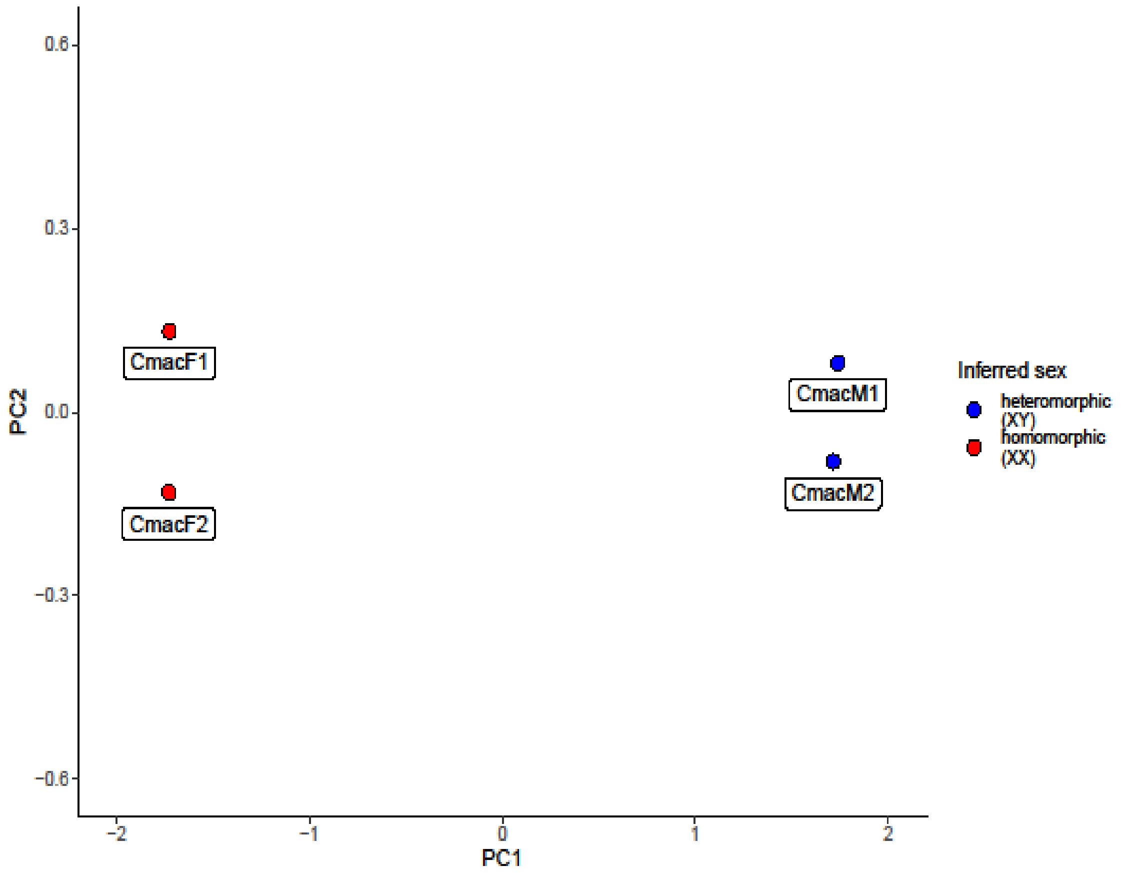

**Fig. SATC|** SATC correctly identifies the sexes of samples.

#### Molecular sexing

To confirm putative Y sequences to be male specific and to establish a protocol for molecular sexing of *C. maculatus* seed beetles, we developed a multiplexed PCR protocol, combining a primer pair for a male specific sequence with a primer pair for an autosomal sequence as a positive control.

##### Primer Design

*Positive control primer pair:*

As the autosomal positive control, we considered known single copy BUSCO genes in the *C. maculatus* genome and scanned for viable primers via FastPCR software [27]. We ended up using the following primer pair (mo25_2F22_2 (5'-CGAGCCTCAGGCTGATATAATTGTAGCAC) & mo25_2R8_6 (5'-GTGCAAATATATTCCACAGTCGGAGACCTA)), amplifying 189bp on the autosomal contig utg000177l_1:

gtgcaaatatattccacagtcggagacctagtcccaatctgccttcgtaaaatattgttaaaaacctgggcaacatccttctttccttcaaaatcaattc
gatttaaattctgtatcagcaataacagtaaattgctattatacaattcttgggctaactgtgctacaattatatcagcctgaggctc

*Y-specific primer pair:*

As a target for the Y specific primer we chose the longest YS contig (utg000322l_1). To identify primers specific for this contig, we split the utg000322l_1 contig into 1kb pieces and blasted the 1kb regions against our reference genome identifying regions that uniquely mapped to utg00322l_1. We then designed the following Y-specific (utg000322l_1-specific) primer pair with FastPCR (utg322_C_1F56_2 (5'-TCAAACATTCCCTTGGAGCTCGTTTGAA) & utg322_C_1R2_2 (5'-ACTGAGGTAGTTCCCCATTTCATGTTACAA)) that amplifies a 297bp on contig utg000322l_1:

tcaaacattcccttggagctcgtttgaaacctctacaattttggtatttaatttcaaacgatggtagagtatctacggctcattgtacttgcatggctagaaacagtgaagtttgtagtc
ataatgtggcatgattccatgcttgaatactgcacataatgcaaatagaaaaaaagaagataagcaagaaggatgttatgtaccgattgatatgcttgcagagtggccaaaggca
aattcatctacctaagtgccaattaagcctatttgtaacatgaaatggggaactacctcag

*PCR protocol*

We optimized the multiplexed PCR protocol to achieve reliable sexing and ended up with the following protocol. Note that the autosomal primer target is diploid while the Y-specific sequence in males is haploid, which is reflected in the primer concentration with the aim to have both primer products to be similarly detectable on a gel. We used virgin adult males and females from the Lomé base population to test the primers.

| Reagents | Concentration in reaction | Composition of cycles | | |
|---|---|---|---|---|
| ddH2O | | **Stage** | **°C** | **Time** |
| Taq buffer | 1X | Pro-incubation | 94 | 5min |
| MgCl2 | 2.5 mM | Denaturation | 94 | 30sec |
| mo25_2F22_2 | 0.4 µM | Annealing | 64 | 30sec |
| mo25_2R8_6 | 0.4 µM | Extension | 72 | 30sec |
| utg322_C_1F56_2 | 1 µM | Final Extension | 72 | 10min |
| utg322_C_1R2_2 | 1 µM | Standby temperature | 4 | |
| dNTPs | 0.4 mM | Number of cycles | 35 | |
| Taq polymerase | 0.025 u/µL | | | |
